# Opposing chromatin remodelers control transcription initiation frequency and start site selection

**DOI:** 10.1101/592816

**Authors:** Slawomir Kubik, Drice Challal, Maria Jessica Bruzzone, René Dreos, Stefano Mattarocci, Philipp Bucher, Domenico Libri, David Shore

## Abstract

Precise nucleosome organization at eukaryotic promoters is thought to be generated by multiple chromatin remodeler (CR) enzymes and to affect transcription initiation. Using an integrated analysis of chromatin remodeler binding and nucleosome displacement activity following rapid remodeler depletion, we investigate the interplay between these enzymes and their impact on transcription in budding yeast. We show that many promoters are acted upon by multiple CRs that operate either cooperatively or in opposition to position the key transcription start site-associated +1 nucleosome. Functional assays suggest that +1 nucleosome positioning often reflects a trade-off between maximizing RNA Polymerase II recruitment and minimizing transcription initiation at incorrect sites. Finally, we show that nucleosome movement following CR inactivation usually results from the activity of another CR and that in the absence of any remodeling activity +1 nucleosomes maintain their positions. Our results provide a detailed picture of fundamental mechanisms linking promoter nucleosome architecture to transcription initiation.

## Introduction

The availability of information encoded in eukaryotic genomes is restricted by wrapping of the DNA helix in nucleosomes, the basic units of chromatin. Regulation of the accessibility of chromosomal DNA to transcription factors (TFs) and the transcriptional machinery itself is believed to play an important role in keeping certain genes silenced while permitting transcription initiation at precisely defined positions at active genes. Such tight and selective regulation is reflected by canonical patterns of promoter nucleosome architecture typically followed by phased arrays of nucleosomes over the downstream gene bodies (Lai and Pugh, 2017). Thus, at promoters of active genes, the transcription start site (TSS) is typically located just upstream of the dyad axis of a well-positioned nucleosome termed “+1” which is followed by regularly spaced genic nucleosomes (+2, +3, etc.). Directly upstream of the +1 nucleosome one typically finds a region of accessible chromatin, termed the nucleosome-depleted region (NDR), whose size is gene-dependent and which at some promoters can be occupied by an unstable nucleosome-like particle termed a “fragile nucleosome” (FN) (Brahma and Henikoff, 2018; Henikoff et al., 2011; Kent et al., 2011; Kubik et al., 2015; Weiner et al., 2010; Xi et al., 2011). The position of the +1 nucleosome and the existence of the NDR are crucial for recruitment of the transcriptional machinery and initiation of transcription (Kubik et al., 2018; Rhee and Pugh, 2012).

ATP-dependent chromatin remodelers (CRs), multi-subunit molecular machines that utilize the energy of ATP hydrolysis to slide, eject or modify nucleosomes (Clapier et al., 2017), have emerged as major factors shaping the chromatin landscape at promoters (Lai and Pugh, 2017). There are four main CR subfamilies – SWI/SNF, ISWI, CHD and INO80 – all of which are conserved from yeast to humans. Each subfamily displays unique biochemical activities and is associated with specific roles in the cell (Clapier et al., 2017). SWI/SNF subfamily members, in budding yeast represented by the essential remodeler RSC and the canonical SWI/SNF remodeler, slide nucleosomes to the edge of linear DNA templates in vitro, maximizing the amount of nucleosome-free DNA (Flaus and Owen-Hughes, 2003; Kassabov et al., 2003). They are also able to displace histone octamers from the DNA template (Boeger et al., 2003; Clapier et al., 2016; Lorch et al., 2011; Lorch et al., 2006). In vivo, RSC and SWI/SNF participate in the generation of NDRs at promoters by at least two mechanisms: sliding the +1 and -1 nucleosomes away from each other (Badis et al., 2008; Ganguli et al., 2014; Krietenstein et al., 2016; Kubik et al., 2015; Yen et al., 2012) and destabilizing or ejecting promoter nucleosomes (Boeger et al., 2003; Floer et al., 2010; Klein-Brill et al., 2019; Kubik et al., 2015). ISWI type (represented by ISW1 and ISW2 in budding yeast) and CHD type (CHD1) CRs equalize the length of the naked DNA on both sides of a nucleosome in vitro (Stockdale et al., 2006). In vivo they have a predominant role in setting the spacing of intragenic nucleosomes, with CHD1 acting mostly at genes with shorter linkers than ISWI (Gkikopoulos et al., 2011; Ocampo et al., 2016; Tirosh et al., 2010; Whitehouse et al., 2007). Curiously, despite acting mostly within gene bodies, binding by these CRs is detected predominantly at gene promoters (Zentner et al., 2013). Finally, members of the INO80 family, SWR-C and INO80, are implicated in deposition and removal of histone H2A.Z, respectively (Kobor et al., 2004; Krogan et al., 2003; Mizuguchi et al., 2004; Papamichos-Chronakis et al., 2011), although the latter function is currently controversial (Wang et al., 2016; Watanabe and Peterson, 2016). In vitro, INO80 but not SWR-C can slide nucleosomes similarly to ISWI (Udugama et al., 2011) and recently was also shown to be able to move a significant number of +1 nucleosomes to in vivo-like positions on a reconstituted yeast chromatin template (Krietenstein et al., 2016). The actual in vivo chromatin state likely results from the interplay between these diverse remodeling activities but the links between them are just starting to emerge.

Several studies, in both yeast and mammalian cells, point to cooperative or opposing activities of certain CRs (Gkikopoulos et al., 2011; Mohd-Sarip et al., 2017; Morris et al., 2014; Parnell et al., 2015; Rawal et al., 2018; Tomar et al., 2009; Yen et al., 2012). It is thus of interest to learn how the activities of the complete set of these enzymes are intertwined. Since RSC is the only yeast CR essential for cell viability, the activities of all other CRs in live cells have been studied predominantly by the use of deletion mutants. However, this approach carries the risk of underestimating the role of individual CRs due to compensating effects (El-Brolosy and Stainier, 2017). Recent analysis with in vitro-reconstituted chromatin and purified remodeling complexes (Krietenstein et al., 2016) suggests that nucleosome positioning in live cells is achieved by the combined action of CRs and specific transcription factors. However, since many cellular processes such as transcription or replication were not reconstituted in vitro, and the concentrations of proteins used do not necessarily reflect the physiological state, this model awaits rigorous testing in vivo.

By integrating the analysis of novel remodeler binding data with nucleosome occupancy and position changes upon conditional depletion of these complexes we obtained insights into their functionality and the interplay between them. We show that promoter nucleosome arrangements are the net result of combined activities of collaborating and opposing CRs. We demonstrate that the majority of nucleosome rearrangements observed in the absence of a remodeler are caused by the antagonizing activity of other enzymes. As a consequence, the in vivo position of +1 nucleosomes is often determined by the activities of two opposing groups – “pushers” and “pullers” – and has a significant effect on both RNA polymerase II (RNAPII) initiation rates and TSS selection. Remarkably, removal of all “pushers” and “pullers” leaves +1 nucleosome positions largely unaffected, in contrast to removal of only one type of remodeler. Our results provide a detailed picture of mechanisms leading to the establishment of promoter nucleosome architecture and the functional significance of +1 nucleosome position.

## Results

### Chromatin remodelers bind intergenic regions in specific combinations

To investigate the links between the activities of different CRs we performed parallel measurements of remodeler binding, using chromatin endogenous cleavage (ChEC)-seq (Zentner et al., 2015) and nucleosome occupancy changes upon conditional depletion of each remodeler, by MNase-seq (**Figure 1A**). ChEC-seq signals for individual remodeler subunits (Rsc8, Swi3, Isw2, Ino80, Isw1, Chd1) were normalized to the signal obtained in a strain expressing “free” MNase under the control of the *REB1* promoter (see Methods for details). Since SWR-C is known to lack nucleosome sliding activity (Udugama et al., 2011) we did not investigate this remodeling complex. A distinct binding pattern was observed for each remodeler, the common point being that signal peaks were found predominantly in intergenic regions, with the majority located at gene promoters (**Figure 1B and S1A**). Rsc8 binds at a large number of gene promoters whereas Swi3, Isw2 and Ino80 bind with similar strength at a much smaller subset (**Figure S1B**). Isw1 and Chd1 displayed low binding at a large number of promoters and a higher signal within the coding regions than other CRs. Due to a frequent overlap between regions bound by various CRs we decided to assemble a single common list of all regions bound by at least one remodeler (see Methods) and calculated normalized ChEC signal of every remodeler in each of these regions. By comparing binding signals of all CRs, we found that RSC and SWI/SNF bind mostly exclusively and have distinct subsets of targets (**Figure S1C**), consistent with previous observations (Yen et al., 2012). Moreover, the binding signal of Swi3 was strongly anti-correlated with that of Isw1 and Chd1 and there was also a significant correlation between the signal of both Isw2 and Ino80 as well as between Isw1 and Chd1 (**Figure S1D**).

**Figure 1.**
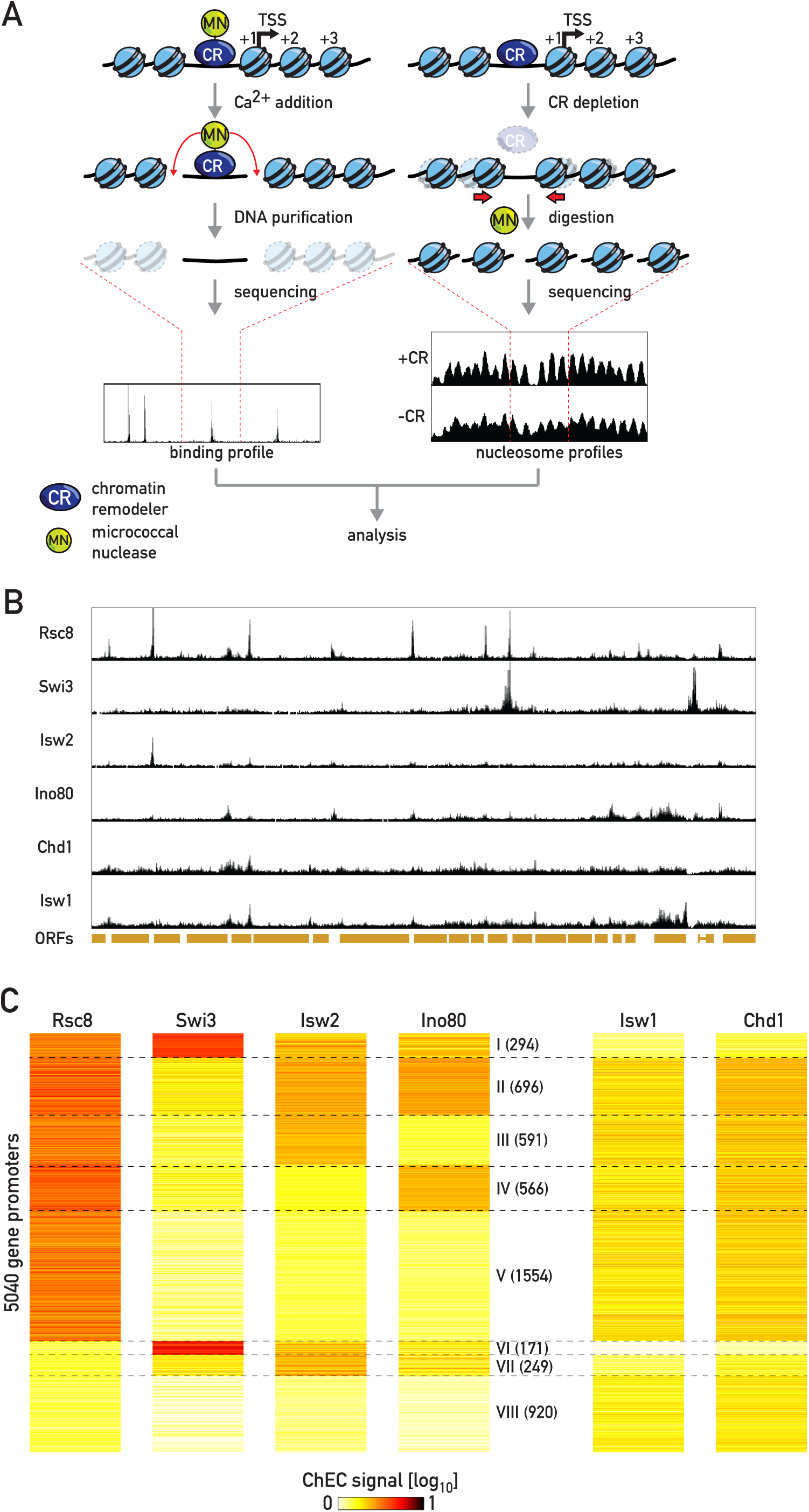
Chromatin remodelers (CRs) bind in defined combinations. (**A**) Schematic representation of the experimental setup. CR binding was measured by ChEC-seq (left). Remodeler activity was evaluated as the local change in nucleosome occupancy measured by MNase-seq (right) following conditional depletion of the catalytic subunit of a CR. (**B**) Snapshot of a genomic region displaying normalized ChEC-seq signal for each CR. (**C**) Heatmap representing normalized remodeler ChEC signal at gene promoters clustered by k-means (k=8).

To investigate remodeler co-occurrence at distinct intergenic regions, including promoters, we performed k-means clustering (k=8) of ∼5000 gene promoters (see Methods) based on the binding signal for CRs known to affect promoter nucleosomes (RSC, SWI/SNF, ISW2, and INO80; **Figure 1C** and **S1E, Table S1**). ISW1 and CHD1 were not included in the clustering as their promoter binding signal was relatively low and these complexes are known to act predominantly in coding regions (Gkikopoulos et al., 2011; Tirosh et al., 2010; Whitehouse et al., 2007). We detected RSC binding at most promoters (cluster I-V) with a small subset of them also bound by SWI/SNF (cluster I). RSC frequently bound with ISW2 and/or INO80 CRs (clusters I-IV) whereas SWI/SNF predominantly associated with ISW2. Cluster V displayed binding by RSC with no other remodeler bound, whereas cluster VII was bound most prominently by ISW2, sometimes together with INO80. Cluster VIII did not show a significant signal for any remodeler. Promoters in different clusters display diverse nucleosome arrangements (**Figure S1F**) with SWI/SNF-bound clusters I and VI having the broadest NDRs and clusters VII-VIII, not bound by either RSC or SWI/SNF, the narrowest. SWI/SNF-bound clusters are also associated with the highest transcription rate while cluster VIII with the lowest (**Figure S1G**). Moreover, SWI/SNF-bound clusters contain promoters bearing the canonical TATA-box more frequently than other clusters (**Fig S1H**). Finally, several clusters are enriched for specific functional categories based upon GO-terms analysis (**Figure S1I**), suggesting that their unique remodeler configurations might play some role in their co-regulation. In summary, clustering analysis points to the existence of a limited number of remodeler combinations present at particular genomic locations.

### Effects of remodeler depletion define three distinct remodeler classes

To reveal how the activities of CRs interact to establish genomic nucleosome patterns we compared nucleosome occupancy changes upon conditional depletion of chromatin remodeler catalytic subunits at different genomic regions. Rapid depletion of chosen proteins was achieved utilizing either the anchor away (Haruki et al., 2008) (for Snf2, Isw2, Chd1 and Swc2) or AID* degron (Morawska and Ulrich, 2013; Nishimura et al., 2009) (for Ino80 and Isw1) systems (**Figures S2A** and **S2B**). For RSC we used our previously published Sth1 depletion data (Kubik et al., 2018).

Each remodeler depletion caused distinct changes to nucleosome occupancy patterns (**Figure 2A**). RSC depletion caused shrinkage of the NDR due to upstream movement of the +1 nucleosome and downstream movement of the -1 (**Figure 2B**; (Badis et al., 2008; Ganguli et al., 2014; Kubik et al., 2015; Parnell et al., 2015; van Bakel et al., 2013)), but also led to stabilization of a subset of FNs (**Figure S2C and S2D**). Similarly, depletion of the catalytic subunit of the related SWI/SNF complex (Snf2) resulted in an upstream +1 nucleosome shift (**Figure 2B**) and stabilization of FNs (**Figure S2C and S2D**) but the number of affected regions was much lower than for RSC. We also note that genes displaying SWI/SNF-mediated nucleosome rearrangements are characterized by an unusually large NDR (**Figure S2E**) often occupied by more than one FN.

**Figure 2.**
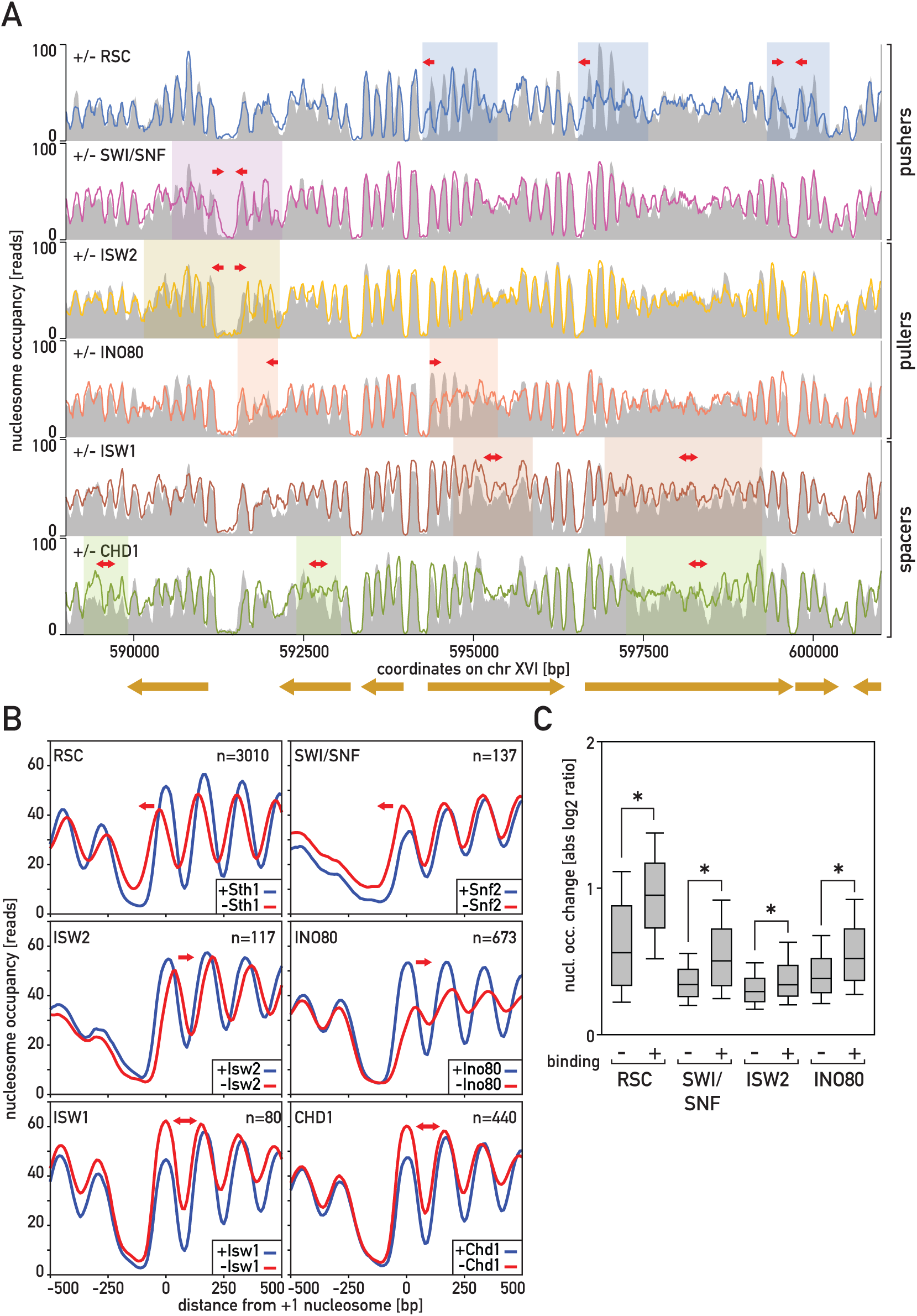
CRs display three broad types of activity. (**A**) Snapshot of a genomic region displaying nucleosome occupancy in wild-type (grey area) or CR-depleted cells (colored lines); red arrows and shaded areas indicate regions and directions of significant rearrangements for each remodeler depletion. (**B**) Nucleosome occupancy at promoters of genes displaying significant changes upon CR depletion in regions centered on the +1 nucleosome dyad. (**C**) Boxplot comparing nucleosome occupancy changes upon depletion of of RSC, SWI/SNF, ISW2 or INO80 at sites displaying low or high ChEC signal of each depleted remodeler; asterisk indicates significant difference (p<0.05, Mann-Whitney test).

Depletion of INO80 and ISW2 both yielded similar +1 nucleosome repositioning downstream (i.e. away from the NDR) at distinct subsets of genes (**Figure 2B**) with INO80 having an effect on a larger set of genes than ISW2. Curiously, a number of stable nucleosomes, some of them +1 nucleosomes of annotated genes (15% for ISW2 and 57% for INO80), became destabilized (i.e. they were not detected at high levels of MNase digestion) in the absence of these CRs (**Figure S2F and S2G**).

Depletion of ISW1 and CHD1 did not result in any notable changes in +1/-1 nucleosome positions (**Figure 2B**). Rather, genes displayed more disordered intragenic nucleosomes (more variation in peak-to-peak distance and decreased peak heights), consistent with previous studies utilizing deletion mutants ((Gkikopoulos et al., 2011); **Figure 2A**). The previously ascribed role of ISW2 in spacing of genic nucleosomes (Gkikopoulos et al., 2011) appears to stem from its role in setting the position of the +1 barrier, since the nucleosomes of genes with an unaffected +1 following ISW2 depletion did not display any change in spacing or peak height (**Figure S2H**, compare with **Figure 2B**).

Based on the observed effects at promoters and gene bodies, we defined three main groups of nucleosome-repositioning complexes: (1) “pushers” (RSC and SWI/SNF) that can shift nucleosomes away from the NDR and are able to destabilize nucleosomes, (2) “pullers” (ISW2 and INO80) that can shift the +1 (and potentially other intragenic nucleosomes) in the direction of the NDR and (3) “spacers” (ISW1 and CHD1) that control the distance between intragenic nucleosomes without affecting +1 nucleosome position.

Next, we examined the extent to which remodeler binding (**Figure 1**) correlates with nucleosome occupancy changes upon remodeler depletion. To this end, we grouped sites displaying strong remodeler binding signal (>3-fold enrichment over background, see Methods) for each remodeler and measured nucleosome occupancy changes in the surrounding regions upon depletion of the remodeler. These values were then compared to occupancy changes at sites where the remodeler signal was low (<1.5-fold enrichment). In each case we observed significantly higher changes in nucleosome occupancy at remodeler-bound regions (**Figure 2C**). For “spacers” we observed no such trend as the effects of their depletion are prominent in gene bodies despite apparent binding of the complexes at promoters (Zentner et al., 2013). Nevertheless, genes whose promoters were strongly bound by the “spacers” displayed better nucleosome phasing in gene bodies than genes displaying a weak signal (**Figure S2I**). However, both types of genes displayed a similar loss of phasing upon depletion of ISW1 or CHD1.

### Cooperative and opposing activities of multiple chromatin remodelers determine nucleosome positions

Existing evidence suggests that the interplay between CRs might include both cooperative and opposing interactions (Morris et al., 2014; Parnell et al., 2015; Rawal et al., 2018; Tomar et al., 2009; Yen et al., 2012). We formulated the following hypotheses which we subsequently tested: (i) CRs with similar activities can act redundantly; (ii) the activity of a remodeler can be suppressed by that of an opposing remodeler(s). The implication of these two propositions would be that the depletion of a remodeler might not yield a measurable effect on nucleosome occupancy/position near its binding site due to compensation by synergistic remodeler(s) or in cases where an opposing remodeler is simultaneously depleted or simply not present. Indeed, nucleosome occupancy changes measured upon depletion of RSC or SWI/SNF at sites strongly bound by both enzymes (see Methods) were weaker than at sites bound by just one of them (**Figure S3A**). Similarly, the effects of ISW2 depletion were stronger at clusters bound by ISW2 only than at clusters bound by ISW2 and INO80 (**Figure S3B**). In contrast, the effects of INO80 depletion were stronger when it bound together with ISW2 rather than alone (**Figure S3C**). However, binding of INO80 was weaker when it bound alone relative to cases where it bound together with ISW2 (**Figure S3D**).

To test CR redundancy more directly we simultaneously depleted both “pushers” or both “pullers” and compared the resulting nucleosome rearrangements to those of the corresponding single depletions. Double depletion of RSC and SWI/SNF led to stronger changes at sites bound by both CRs than either single depletion (**Figure 3A and 3B**), consistent with a recent report (Rawal et al., 2018). Interestingly, RSC – SWI/SNF double depletion also had a stronger effect at sites bound by SWI/SNF but not RSC (cluster VI). This might indicate that upon Snf2 depletion, RSC is recruited to SWI/SNF targets and partially substitutes for this remodeler. We did not observe similar behavior of SWI/SNF at sites bound by RSC alone (clusters III-V). ISW2 - INO80 redundancy was already evident at the level of cell growth: the double depletion is lethal, contrary to either single depletion (**Figure 3C**). Consistent with this finding, in every cluster bound by ISW2 or INO80 the double depletion had stronger effects than either single depletion (**Figure 3D**), leading to widespread aberrations in nucleosome patterns (downstream shifts and/or destabilization of +1 nucleosomes, and impaired phasing in gene bodies) that were qualitatively similar to single deletions but amplified in magnitude and number of affected genes (**Figure 3E and 3F**). Similarly, simultaneous depletion of both “spacers” led to a stronger loss of nucleosome phasing in gene bodies than either single depletion, consistent with observations from the corresponding deletion mutants (Gkikopoulos et al., 2011) (**Figure S3E**). In summary, our analysis indicates a considerable degree of redundancy between RSC and SWI/SNF and between ISW2 and INO80.

**Figure 3.**
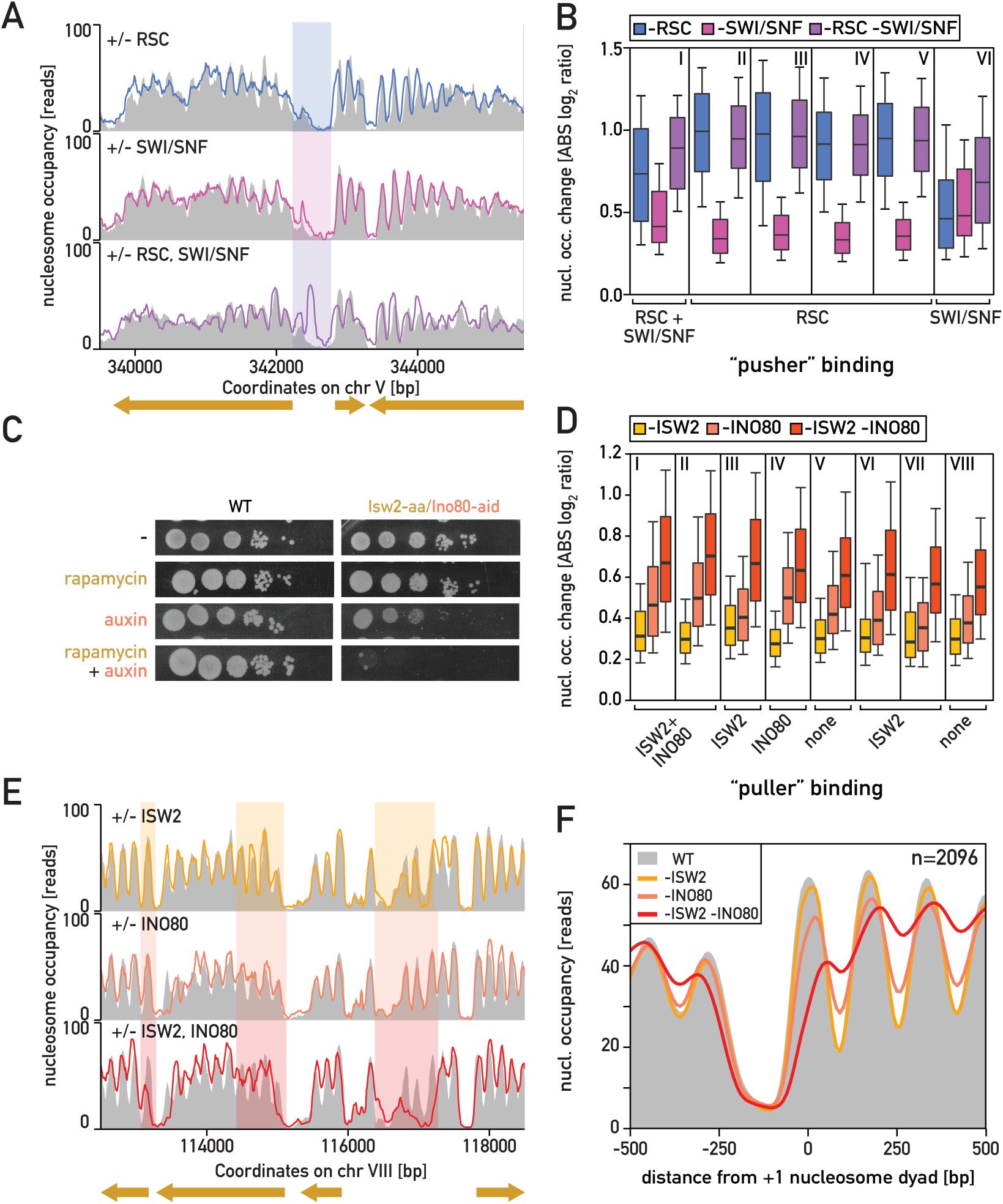
CRs with similar activities act redundantly. (**A**) Snapshot of a sample genomic region displaying stronger nucleosome occupancy change upon simultaneous RSC and SWI/SNF depletion compared to depletion of individual complexes. (**B**) Nucleosome occupancy changes upon depletion of RSC, SWI/SNF or both CRs simultaneously calculated in each cluster binding these complexes. (**C**) Spot assay of wild-type yeast strain (left) and strain in which Isw2 was tagged with FRB in order to deplete it with rapamycin and Ino80 was tagged by aid* for auxin-mediated depletion (right) plated on medium containing rapamycin, auxin or both chemicals simultaneously. (**D**) Nucleosome occupancy changes upon depletion of ISW2, INO80 or both CRs simultaneously calculated for all clusters. (**E**) Snapshot of a representative genomic region displaying a stronger nucleosome occupancy change upon simultaneous ISW2 and INO80 depletion compared to depletion of either individual complex. (**F**) Average nucleosome occupancy plot for wild-type cells and cells depleted of ISW2, INO80 or both CRs simultaneously, averaged over all genes where a significant change of +1 nucleosome occupancy was observed upon simultaneous depletion.

### +1 nucleosomes maintain their positions in the absence of opposing remodeling activities

It is not known what causes the nucleosome repositioning frequently observed upon depletion of a remodeler. Given the opposing activities of “pushers” and “pullers”, we hypothesized that the effects observed upon depletion of one type of remodeler might result from the activity of an enzyme of the other type. Consistent with this idea, the effects of RSC and SWI/SNF depletion were strongest at sites where these CRs co-bound with ISW2 (**Figure S4A**). Conversely, the effects of INO80 depletion were strongest at sites where this remodeler binds alongside both RSC and SWI/SNF compared to sites where it binds with just one “pusher”. Furthermore, the weakest effect of INO80 depletion was observed at sites where neither “pusher” binds nearby (**Figure S4B**).

Our results thus indicate that promoter nucleosome arrangement results from the net activity of specific combinations of CRs and that the absence of a given remodeler might result in nucleosome changes caused by the activity of the remaining remodeling complexes. In order to test this proposition more directly we performed experiments in which we simultaneously depleted “pushers” and “pullers”. Based on our binding data (**Figure 1C**) RSC is often predicted to be opposed by either ISW2 or INO80, whereas SWI/SNF would mostly be counteracted by ISW2. We concentrated on RSC and ISW2/INO80-bound sites due to the high number of regions having these combinations and the fact that depletion of RSC cannot be compensated by SWI/SNF. If nucleosome rearrangements observed upon RSC depletion – upstream shift of the +1 nucleosome and stabilization of FNs – resulted from the activity of ISW2 or INO80 we would expect to observe milder effects upon simultaneous depletion of RSC with ISW2 or INO80. We first measured nucleosome occupancy centered on the +1 nucleosome of genes associated with RSC and ISW2, in a wild-type strain and in strains where one or both complexes were depleted (**Figure 4A**). Interestingly, the nucleosome pattern obtained upon simultaneous depletion of RSC and ISW2 was similar to the one obtained upon depletion of RSC alone (**Figure 4A**; compare blue versus green plots). We performed a similar analysis for RSC- and INO80-bound +1 nucleosomes (**Figure 4B**). Here, the effect of double depletion was slightly weaker than depletion of RSC alone, as the upstream +1 shift was somewhat less pronounced. Still, the +1 nucleosome clearly shifted upstream in the absence of both CRs and the changes upon double depletion generally resembled RSC depletion alone. These observations might result from (i) redundant activity of the “pullers”, (ii) activity of “spacer” CRs that is more pronounced in the absence of “pullers”, or (iii) an inherent preference of nucleosomes for positions that are more upstream than those observed in wild-type cells. To distinguish between these possibilities, we performed experiments in which we simultaneously depleted RSC together with the two opposing CRs and compared the results to RSC alone or ISW2/INO80 double depletion. At promoters bound by RSC and ISW2, or RSC and INO80, we observed only minor changes in +1 nucleosome position in the absence of all three CRs (**Figures 4C 4D**). These observations lead to an important notion, namely that when chromatin remodeling is shut down by multiple CR depletion +1 nucleosomes tend to remain relatively close to their positions under wild-type conditions.

**Figure 4.**
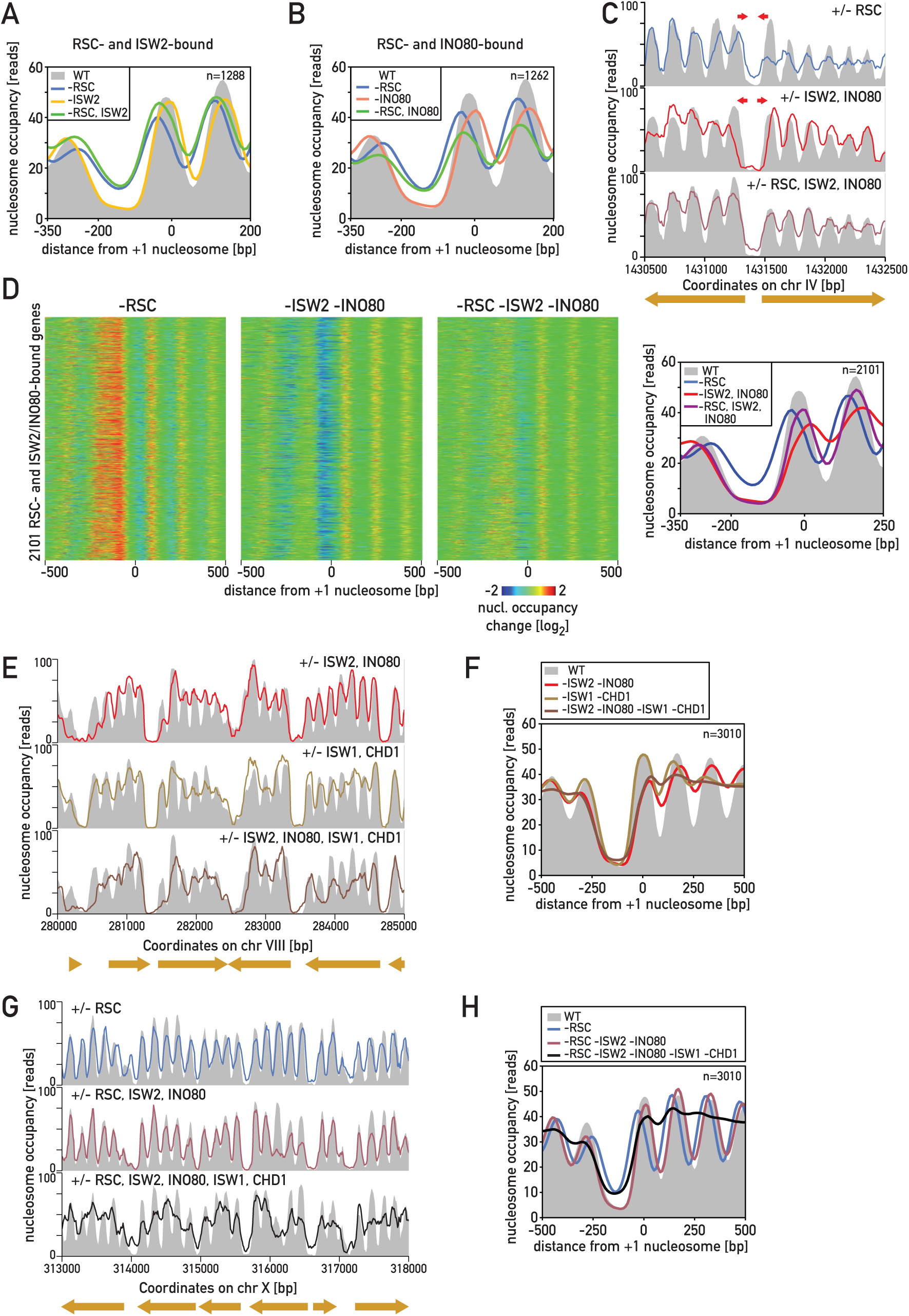
Position of +1 nucleosome results from the net activity of multiple cooperating and opposing CRs. (**A**) Average nucleosome occupancy upon depletion of RSC, ISW2 and both CRs simultaneously at all genes displaying binding of RSC and ISW2. (**B**) As (A) but for RSC and INO80-bound promoters. (**C**) Snapshot of a representative genomic region displaying strong opposing changes in +1 nucleosome position upon depletion of RSC or the two “pullers” (ISW2 and INO80) and only minor changes upon simultaneous depletion of all three CRs. (**D**) Heatmaps of nucleosome occupancy change (first three panels) and average plots of nucleosome occupancy (right panel) for cells depleted of either “pullers” or RSC and “pullers” simultaneously, at genes bound by these CRs. (**E**) Snapshot of a sample genomic region displaying nucleosome occupancy changes shown for depletion of the “pullers” (ISW2, INO80), the “spacers” (ISW1, CHD1) and all four CRs simultaneously. (**F**) Nucleosome occupancy in wild-type cells and cells depleted of both “pullers”, both “spacers” and all four CRs simultaneously, averaged over all genes displaying +1 nucleosome occupancy changes upon RSC depletion. (**G**) Snapshot of a representative genomic region displaying nucleosome occupancy changes following depletion of RSC (top), RSC and “puller” (RSC, ISW2, INO80; middle), and RSC, “puller” and “spacer” simultaneously (bottom). (**H**) Average nucleosome occupancy plots for wild-type cells and cells depleted of RSC, RSC and both “pullers” and RSC, “pullers” and “spacers” simultaneously.

Although ISW1 and CHD1 act predominantly within gene bodies, they could influence the position of the +1 nucleosome in a way that might be masked by other CRs. To test this idea we simultaneously depleted “pullers” and “spacers” and compared the results to depletion of only the two “pullers”. We found that the +1 nucleosome position is very similar under these two conditions, in either the presence of RSC (**Figures 4E and 4F**) or in its absence (**Figures 4G and 4H**), arguing against an additional role for “spacers” in +1 positioning. Nevertheless, genic nucleosome phasing was more strongly disrupted upon depletion of “spacers” and “pullers” compared to “spacers” alone, presumably due to the altered position of the +1 nucleosome, proposed to act as a barrier against which downstream genic nucleosome are phased through a “statistical positioning” mechanism (Hughes and Rando, 2014; Kornberg and Stryer, 1988). In summary, these results indicate that while most +1 nucleosomes remain robustly positioned in the absence of remodeling activity, genic nucleosomes significantly change their positions when “spacers”, or “spacers” plus “pullers” are depleted.

We also noted that nucleosomes which became destabilized upon ISW2 or INO80 depletion (i.e. became FNs; **Figure S2G**) were much less affected upon simultaneous depletion of RSC and both “pullers” (**Figure S4C** and **S4D**). This suggests that their destabilization results from a destructive activity of RSC. Moreover, the FNs that became stable upon RSC depletion were not stabilized when RSC was co-depleted with both “pullers” (**Figure S4E**). Co-depletion of RSC with only one “puller” did not prevent FN stabilization indicating that ISW2 and INO80 are also redundant with respect to this aspect of chromatin organization. Taken together, these results imply that nucleosome destabilization in the absence of “pullers” is due to the destructive activity of RSC. Conversely, substitution of an FN with a stable nucleosome following RSC depletion is mediated by “pullers”.

### Combined remodeler action at +1 nucleosome influences TBP binding and TSS selection

We showed recently that the role of RSC in +1 nucleosome placement is crucial for binding of TBP, a key step in RNA Polymerase II (RNAPII) recruitment (Kubik et al., 2018). However, it is still unknown which steps of gene activation are affected by SWI/SNF, or by the absence of the “pullers”, which should generally increase the accessibility of the TBP binding site. To explore the role of these CRs in gene expression we measured binding of TBP (Spt15) and the RNAPII catalytic subunit (Rpb1) by ChIP-seq following their depletion.

Genes down-regulated in the absence of SWI/SNF (n=128, at least 1.5-fold decrease in RNAPII levels) displayed a prominent decrease in TBP binding at their promoters that was linked to an upstream shift of the +1 nucleosome and an increase in nucleosome occupancy at the TATA element (**Figure 5A**). The genome-wide correlation between TBP and RNAPII occupancy change upon SWI/SNF depletion was strong (**Figure S5A**), consistent with a causal relationship. These results indicate that SWI/SNF, like RSC, facilitates transcription by promoting TBP binding and PIC assembly. Unexpectedly, a small number of genes (n=39) actually displayed an increase in RNAPII association and a modest increase in TBP signal upon SWI/SNF depletion. However, these effects were not associated with significant nucleosome occupancy changes (**Figure S5B**).

**Figure 5.**
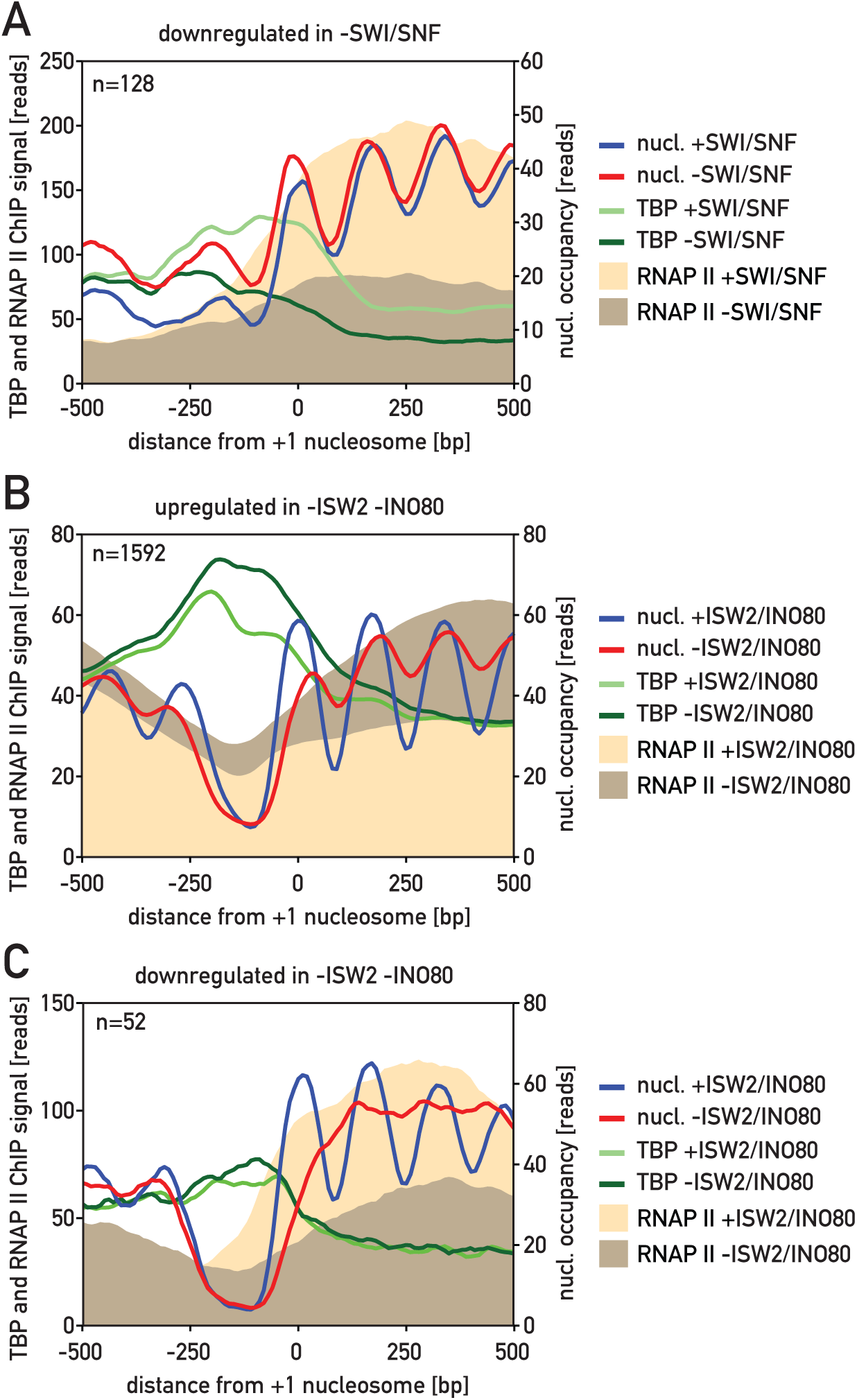
Changes in +1 nucleosome occupancy are linked to transcriptional down- and up-regulation. (**A**) Plots of nucleosome occupancy, RNAPII and TBP ChIP signals, in the presence and absence of SWI/SNF, at genes displaying a significant decrease in RNAPII level upon SWI/SNF depletion. (**B**) Plots of nucleosome occupancy, RNAPII and TBP ChIP signals, in the presence and absence of ISW2 and INO80, at genes displaying a significant increase in RNAPII level upon simultaneous depletion of ISW2 and INO80. (**C**) As (B) but for down-regulated genes.

Consistent with ISW2-INO80 redundancy with respect to +1 positioning, we observed relatively few genes where their single depletion had a pronounced effect on either TBP binding or RNAPII levels (**Figure S5C-E**). As expected, double depletion of these two “puller” CRs produced more significant effects, with many more genes displaying increased RNAPII levels (n=1592, versus 45 and 104 for ISW2 and INO80 single depletions, respectively) and only a few with lower levels (n=52) (**Figure 5B, C**). Nevertheless, and as was the case for INO80 depletion alone, down-regulation upon double depletion was not correlated with a TBP signal change (**Figure S5F**). Consistent with this, “puller” double depletion was associated with downstream movement of +1 nucleosome regardless of the transcriptional outcome (**Figure 5B, C**). Regarding the “spacers”, we found that depletion of ISW1 led to both decreases (n=503) and increases (n=332) in RNAPII levels within gene bodies without any significant changes in +1 nucleosome position or promoter TBP binding yet a modest loss of genic nucleosome positioning (**Figure S5G, H**). In contrast, the absence of CHD1 caused an increase in RNAPII levels at 116 genes accompanied by negligible changes in genic nucleosome positioning and no change in TBP binding (**Figure S5I**). No genes displayed a significant decrease in RNAPII levels upon CHD1 depletion.

### “Puller” depletion affects TSS selection by facilitating TBP binding at cryptic downstream TATA elements

In addition to affecting RNAPII initiation rates, nucleosome repositioning can also influence TSS selection at individual genes (Challal et al., 2018; Dreos et al., 2016), possibly giving rise to non-coding transcription (Whitehouse et al., 2007) or altered levels of potentially function transcripts. In order to test how CRs affect transcription initiation events we first performed a genome-wide rapid amplification of 5’ cDNA ends (5’-RACE) analysis (Malabat et al., 2015) in strains depleted for the “pusher” CRs. Depletion of RSC resulted predominantly in an upstream shift of the +1 nucleosome and a decrease of initiation events at genes, as expected ((Ganguli et al., 2014; Kubik et al., 2015; Kubik et al., 2018); **Figure 6A, B**), but no marked change in TSS selection. Similarly, SWI/SNF depletion often led to a decrease in transcription initiation at genes where nucleosomes were rearranged (**Figure 6C, D**). Four genes displayed additional strong signals 3’ of the annotated +1 nucleosome (visible as prominent peaks in the average plot) which were also suppressed in the absence of SWI/SNF (**Figure 6D**). In rare cases, though, RSC or SWI/SNF depletion led to either a shift in the TSS or a change in TSS distribution in situations where two or more predominant initiation sites were observed (**Figure S6A** and **S6B**).

**Figure 6.**
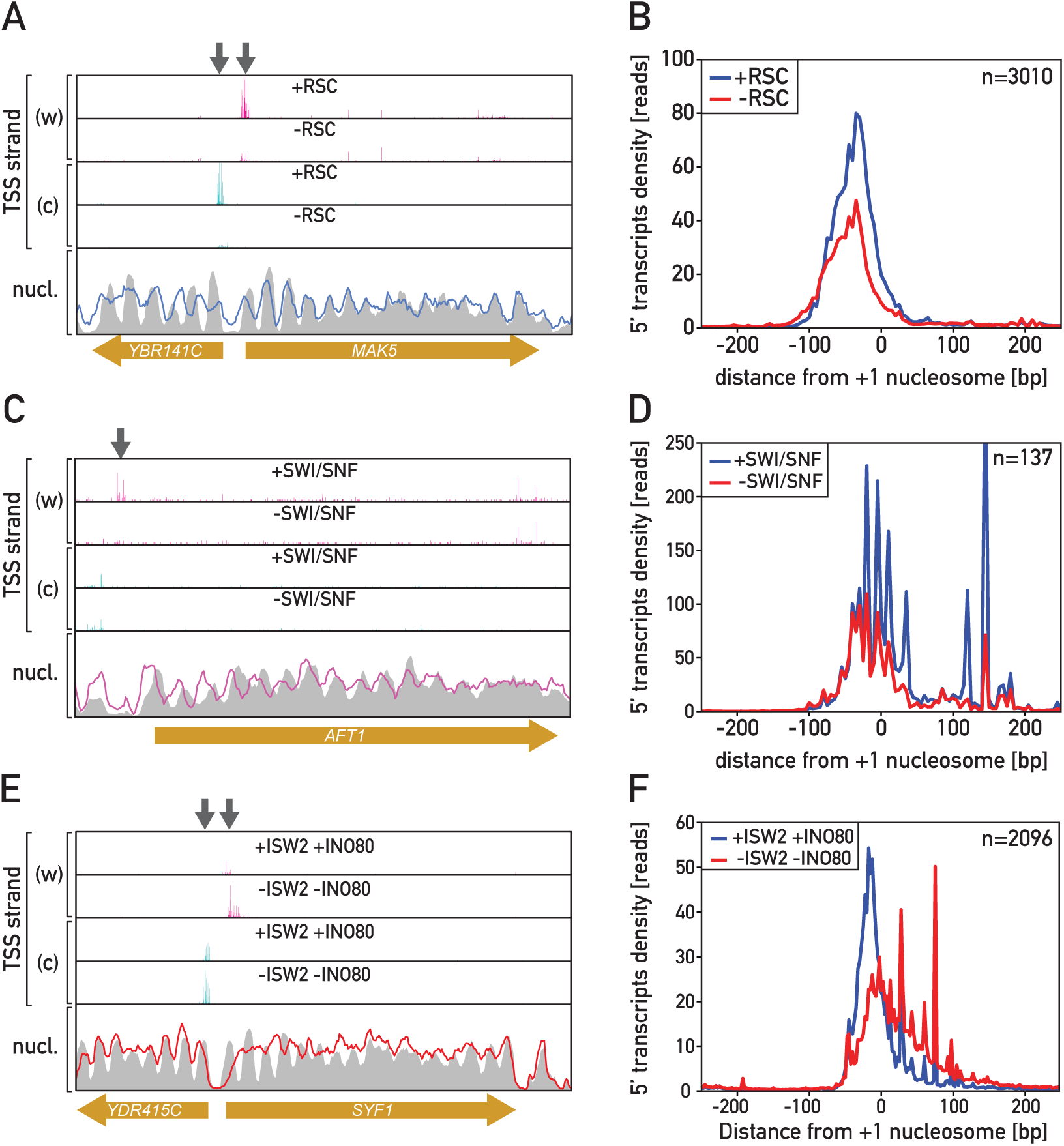
+1 nucleosome shift interferes with transcription start site selection. (**A**) Snapshot of genomic region showing 5’RACE signal (“TSS”) for the Watson (w) and the Crick (c) strands as well as nucleosome occupancy in the presence (grey background) and absence of RSC (colored line). (**B**) Average plot showing 5’RACE signal, in the presence and absence of RSC, for all genes displaying significant occupancy changes at their +1 nucleosome upon RSC depletion. (**C**) As in (A) but for depletion of SWI/SNF. (**D**) As in (B) but for depletion of SWI/SNF. (**E**) As in (A) but for simultaneous depletion of ISW2 and INO80. (**F**) As in (B) but for simultaneous depletion of ISW2 and INO80.

Notably, simultaneous “puller” depletion had a dramatic effect on TSS selection at a large number of genes, often leading to a decrease of initiation at the wild type TSS and the appearance of high levels of novel initiation events located downstream (**Figure 6E, F** and **Figure S6C**). When we plotted 5’-RACE signals at genes with increased RNAPII levels upon “puller” depletion we often observed increases in the signal downstream of the TSS without prominent decreases at the original TSS position (**Figure S6D**), which was not evident at those genes where transcription decreased **Figure S6E**).

To investigate in more detail possible links between +1 nucleosome shifts and the appearance of novel TSSs we first identified the single, strongest TSS at each gene following ISW2/INO80 co-depletion and retained those having a normalized signal of at least 150 reads (n=3524 genes). We then measured signal at these TSSs in wild-type conditions and sorted all genes according to the ratio of the “puller” double-depletion to wild-type signal. This identified a large number of sites (n=1372) where transcription clearly initiated more frequently following “puller” depletion (**Figure 7A**, “increase”, top) but also a smaller number of sites (n=377) where the initiation events became less frequent (“decrease”, bottom). Most of the genes with up-regulated novel start sites have well-annotated TSSs in wild-type cells (n=1210 out of 1372; (van Bakel et al., 2013), and in most cases (n=946) the novel prominent TSSs following “puller” double depletion were more than 20 bp downstream from the wild-type TSS (**Figure 7B**). In only 73 cases was the novel TSS more than 20 bp upstream from the wild-type site. Nevertheless, both cases were associated with very similar downstream shifts of the +1 nucleosome (**Figure 7C** and **Figure S7**).

**Figure 7.**
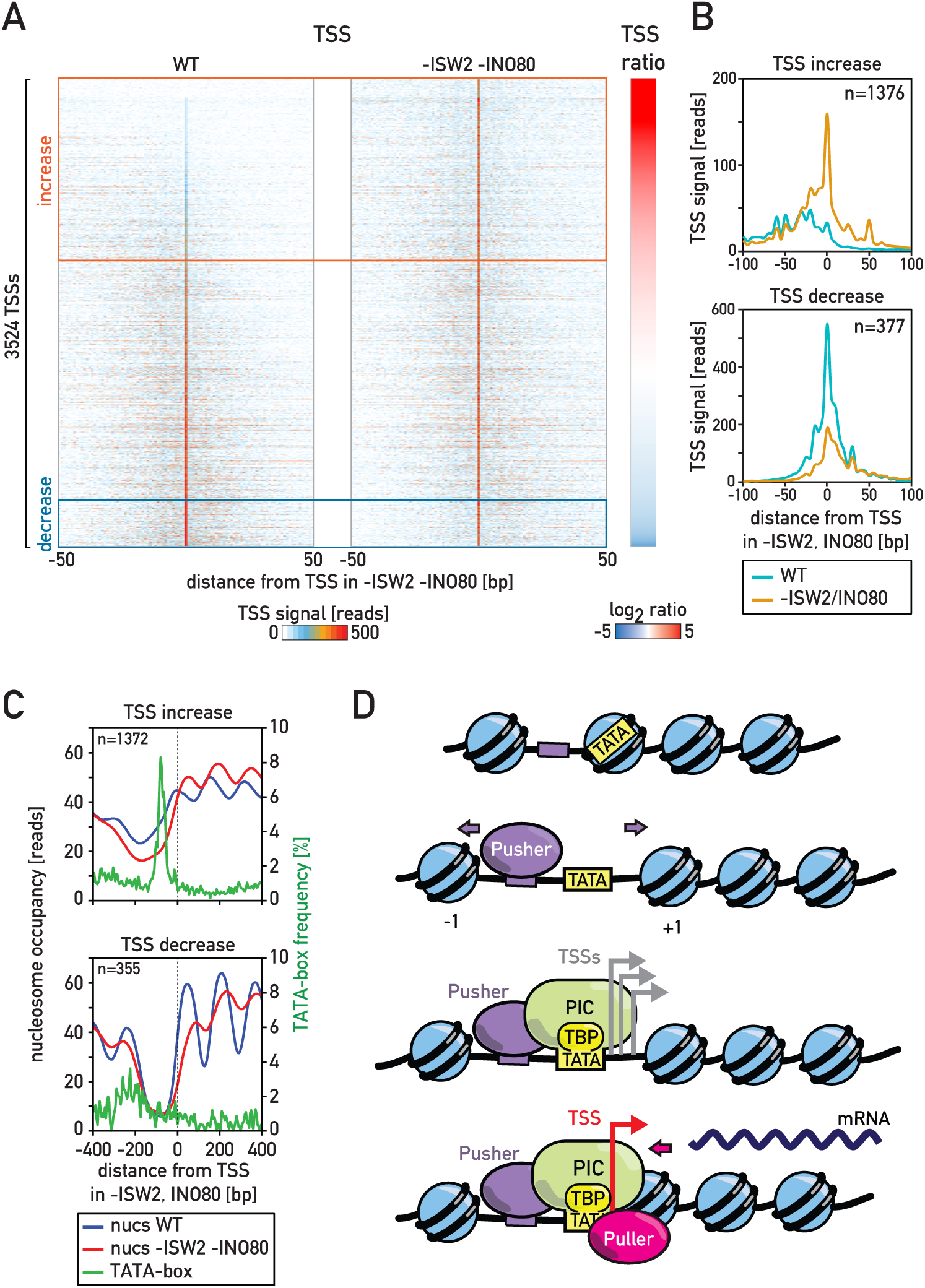
Enhance of TSS usage depends on the presence of TATA-box. (**A**) Heatmap showing 5’RACE signal in wild-type cells (left) and cells depleted of ISW2 and INO80 centered on predominant TSS site determined in the absence of these CRs for each gene (**B**) Average 5’RACE signal for genes displaying most significant increase (top) or decrease (bottom) in the signal. (**C**) Plots displaying nucleosome occupancy in the presence (blue) and absence (red) of ISW2 and INO80 as well as average frequency of the consensus TATA-box motif (green) for genes displaying most significant increase (top) or decrease (bottom) in the 5’RACE signal. (**D**) Schematic representation of mechanisms determining +1 nucleosome position and TSS selection at active genes. Recruitment of “pushers” such as RSC might be guided by specific DNA motifs or TFs leading to creation/expansion of the NDR, exposition of TBP binding sites (TATA) and formation of the PIC; “pullers” reposition the +1 nucleosome to reduce NDR size and to restrict transcription initiation to the position observed in wild-type cells.

The observations outlined above begged the question of why genes with similar nucleosome rearrangements following “puller” depletion would vary so much in the intensity and position of TSS changes. The likely answer became apparent when we plotted the distribution of TATA-box motifs for both up-regulated and down-regulated genes (**Figure 7C**). At sites where transcription initiated more frequently after “puller” depletion we often found a canonical TATA-box peak within 150 bp of the strongest new TSS (n=661 genes), at a position where +1 nucleosome occupancy decreased following “puller” depletion. This was not the case at genes where a decrease in initiation frequency was observed, suggesting that increased downstream initiation following “puller” depletion is often due to the exposure of a “cryptic” TATA box normally occluded in wild-type cells, at least under the conditions of growth employed here. This interpretation is consistent with the increased downstream TBP binding observed at these genes upon ISW2/INO80 double depletion (**Figure 5B**). These findings indicate that ISW2 and INO80 act together to repress transcription at a large number of genes through a mechanism linked to upstream movement of the +1 nucleosome and occlusion of a TATA element otherwise capable of driving PIC assembly and transcription.

## Discussion

### Cooperative and opposing remodeler activities establish promoter nucleosome landscapes

We describe here a comprehensive examination of chromatin remodeler binding and action at promoters in living cells. Our approach of measuring nucleosome positions by MNase-seq immediately following rapid depletion of individual or multiple CRs is likely to reveal the direct action of these enzymes, while avoiding secondary effects that might arise from transcriptional changes that occur in gene deletion strains. Comparing nucleosome occupancy changes following remodeler depletion to ChEC analysis of remodeler localization provides additional insights into remodeler targeting and redundancy. Our results imply a central role for CRs in determining transcription initiation rates but also reveal an unanticipated role for these factors in determining precisely where transcription starts at individual genes.

Our findings establish two different remodeler groups with respect to positioning of the canonical +1/-1 promoter nucleosomes, which we refer to as the “pushers” and “pullers”, the former acting to expand the NDR, the latter to contract it. We identify RSC as the most pervasive promoter-directed remodeler, acting as a “pusher” at a majority of protein-coding genes, consistent with previous reports (Badis et al., 2008; Ganguli et al., 2014; Hartley and Madhani, 2009; Klein-Brill et al., 2019; Kubik et al., 2015; Kubik et al., 2018; Parnell et al., 2008; Parnell et al., 2015). We show that the second “pusher”, SWI/SNF, has a much more restricted set of target genes, primarily working in conjunction with RSC at a set of highly-transcribed genes (Rawal et al., 2018), but also alone at a smaller set of stress responsive genes. Conversely, both ISW2 and INO80 function as promoter-specific “pullers” that act to reduce the NDR due to movement of the +1 or -1 nucleosome (or both) towards its center, consistent with previous work (Klein-Brill et al., 2019; Krietenstein et al., 2016; Tomar et al., 2009; Whitehouse et al., 2007; Whitehouse and Tsukiyama, 2006).

A picture that emerges from our study is that of a competition between “pushers” (RSC and SWI/SNF) and “pullers” (ISW2 and INO80), the result of which leads to precise positioning the +1 nucleosome (**Figure 7D)**. This insight followed from our ability to simultaneously deplete cells of both ISW2 and INO80. Unexpectedly, we found that these two CRs are largely redundant at a significant number of genes. Thus, either ISW2 or INO80 alone are largely sufficient to counteract the effect of the “pushers”, whereas cells depleted of both “pullers” display a significant broadening of the NDR. Our results thus suggest that these two CRs probably constitute the main force counteracting the destructive effect of RSC on the nucleosomes flanking NDRs.

We imagine that both “pushers” and “pullers” act immediately following replication fork passage to re-establish promoter nucleosome architecture, which in yeast appears to occur in a matter of minutes (Fennessy and Owen-Hughes, 2016; Vasseur et al., 2016; Yadav and Whitehouse, 2016); reviewed in (Ramachandran et al., 2017)). Since our studies were carried out on populations of unsynchronized cells and show that depletion of one remodeler often leads to nucleosome movement dependent upon another, they suggest that opposing CRs might be continuously acting upon promoter nucleosomes, thus maintaining+1 nucleosome position within a highly limited range. This “spring trap” state appears to play a key role in determining the probability of TBP binding and PIC assembly and may poise the +1 nucleosome to change its position whenever an additional factor (e.g. TF binding) shifts the balance in favor of one of the two opposing activities. One example of such dynamic remodeler regulation could be the +1 nucleosome movement that occurs at hundreds of genes during cell state changes in the yeast metabolic cycle (Nocetti and Whitehouse, 2016).

Our finding that +1 and -1 nucleosomes remain relatively stable upon simultaneous depletion of both “pusher” and both “puller” CRs suggests that neither thermal motion nor some additional active process is sufficient to alter the preferred positioning of these nucleosomes. We imagine that under these conditions relatively few cells passage through S phase, and that the few that do might be largely responsible for the minimal displacements observed. Interestingly, genic nucleosomes, which are likely to be subjected to disruption by RNAPII, display a massive loss of positioning upon multiple CR depletion.

### Global and local control of remodeler action

As pointed out previously (Yen et al., 2012), detection of CR binding by ChIP is notoriously problematic, making it difficult to determine the extent to which these factors are targeted to specific promoters. We showed here that the application of ChEC-seq to remodeler subunits reveals patterns of remodeler binding specificity that correlate well with nucleosome occupancy changes caused by depletion of the corresponding remodeler. Interestingly, these correlations are stronger for the “pushers” (RSC and SWI/SNF) than it is for the “pullers” (INO80 and ISW2), which, despite their limited binding overlap, are often functionally redundant. This discrepancy between INO80 and ISW2 binding and action may indicate that one factor can rapidly replace the other when it is depleted. Alternatively, ChEC may not capture certain functional remodeler-promoter interactions, perhaps due to either their short half-life or because of limited MNase access upon remodeler binding at certain promoters. Finally, we note that RSC can often substitute for SWI/SNF when the latter is depleted, whereas the converse is not the case. This may reflect more stringent co-factor requirements for SWI/SNF binding or activity.

The “pushers” were shown to slide nucleosomes off the edge of DNA templates in vitro, in effect maximizing the size of the linker between nucleosomes (Flaus and Owen-Hughes, 2003; Kassabov et al., 2003). The same principle seems to apply at NDRs in living cells – RSC and SWI/SNF act to increase the length of linker DNA separating the +1 and -1 nucleosomes. Although the precise mechanism(s) by which these two CRs act is still unclear, it is important to note that both RSC and SWI/SNF appear to engulf the nucleosomes upon which they act (Chaban et al., 2008; Dechassa et al., 2008). Given that these CRs move the +1 and -1 nucleosomes in different directions, it would seem likely that they orient their direction of action with respect to some landmark feature(s) of the NDR that separates these nucleosomes. This could result from the inherent length of relatively nucleosome-free DNA in this region due to the presence of nucleosome-disfavoring poly(dA:dT) tracts or the binding of TFs.

The remaining CRs that we examined (ISW2, INO80, ISW1 and CHD1) all possess similar activities in vitro – they slide nucleosomes towards the central position on a DNA template, thus equilibrating linker length on both sides of a core particle (Stockdale et al., 2006; Udugama et al., 2011). However, their in vivo roles vary, with “spacers” exclusively affecting genic nucleosomes (+2, +3, etc.) and “pullers” acting on the +1 nucleosome (**Figure 2**). What could explain this dichotomy? ISW2 and INO80 form larger complexes than ISW1 and CHD1 which might make it more difficult for them to act on densely packed genic nucleosome arrays due to steric hindrance. Moreover, accumulating evidence suggests that “pullers” are targeted to promoters through direct interactions with general or more gene-specific TFs (Bowman and McKnight, 2017), though the only well-documented example to date is that of ISW2 recruitment by Ume6 (Goldmark et al., 2000). Interestingly, though neither ISW1 nor CHD1 are known to associate with promoter-specific factors, fusing CHD1 to the DNA-binding domain of Ume6 leads to nucleosome repositioning at Ume6 binding sites qualitatively similar to that normally carried out by ISW2 (McKnight et al., 2016). Taken together, these findings suggest that promoter-targeted “pullers” recognize NDRs as extremely long linker DNAs of the +1 or -1 nucleosome, which they shorten, whereas “spacers” primarily scan genic regions where they act to equalize linker lengths.

We also observed nucleosome destabilization upon depletion of the “pullers” caused by the destructive activity of RSC or SWI/SNF. This might result from the collision of a mobilized “pusher”-bound nucleosome with an adjacent core particle as shown by in vitro studies of SWI/SNF (Dechassa et al., 2010; Engeholm et al., 2009). “Pullers” might act to directly protect the vulnerable particle or to mediate its re-deposition.

### Significance of +1 nucleosome positioning for transcription

We have shown recently that RSC facilitates gene transcription by globally increasing the accessibility of TBP binding sites (Kubik et al., 2018). Data presented here show that the related chromatin remodeler SWI/SNF has a similar role but that its action is limited predominantly to genes possessing a canonical TATA box in their promoter. The increase in promoter nucleosome occupancy observed in the absence of RSC and SWI/SNF leads to impaired transcription initiation events which become either less frequent or occur at altered positions. Therefore, the “pushers” not only create a “landing spot” for the transcriptional machinery by generating wide NDRs but also participate in accurate TSS selection, consistent with a recent report (Klein-Brill et al., 2019). Interestingly, general regulatory factors known to influence promoter nucleosome occupancy (e.g. Rap1, Abf1 and Reb1; (Ganguli et al., 2014; Kubik et al., 2015; Kubik et al., 2018)) have also recently been shown to suppress spurious initiation events (Challal et al., 2018; Wu et al., 2018).

Previous studies showed that NDR expansion in the absence of ISW2 leads to an increase of ncRNA synthesis (Whitehouse et al., 2007; Yadon et al., 2010). We expand this picture considerably by showing that “puller” double depletion causes widespread activation of novel downstream TSSs, most likely driven by cryptic TATA elements that become functional upon +1 nucleosome re-positioning. Importantly, these novel TSSs are likely to produce functional transcripts in many cases, suggesting that ISW2/INO80 may have a regulatory role.

In conclusion, we provide a comprehensive view of the effect of CRs on promoter nucleosome positioning in a simple eukaryote (budding yeast). Our results reveal a complex interplay between these factors that plays an important role in determining not only transcription initiation rates but also the precise site of initiation. Results and methods established here will provide a basis for future studies to explore the role of CRs in controlling gene expression under variable growth conditions. Finally, since the CRs as well as the general features of promoter nucleosome organization are highly conserved in metazoans, we anticipate that our general findings will be relevant to unraveling promoter function in these more complex systems.

## Acknowledgments

We thank Mylène Docquier and the iGE3 Genomics Platform (https://ige3.genomics.unige.ch/) at the University of Geneva for high throughput sequencing services, Nicolas Roggli for expert assistance with data presentation and artwork, and all members of the Shore lab for comments and discussions throughout the course of this work. M.J.B. was supported in part by an iGE3 Ph.D. student fellowship. D.S. acknowledges funding from the Swiss National Science Foundation (grant no. 31003A_170153) and the Republic and Canton of Geneva.

## Author Contributions

Conceptualization – S.K., D.C., D.L. and D.S.; Formal Analysis – S.K., D.C., R.D., P.B. and D.L.; Investigation – S.K., D.C., S.M. and M.J.B.; Data Curation – S.K., D.C., and R.D.; Writing – Original Draft, S.K. and D.S.; Funding Acquisition – D.S., D.L. and P.B.; Resources – D.S. and D.L.; Supervision, D.S., D.L., and P.B.

## Methods

### Yeast strains

All experiments presented in this study were performed using budding yeast *Saccharomyces cerevisiae* as the model system. Yeast strains used in this study are listed in **Table S2**. For ChIP-seq experiments of Rpb1 addition of a small fraction of crosslinked chromatin obtained from fission yeast *Schizosaccharomyces pombe* was used as a spike-in control. Cells were grown in YPAD medium at 30°C for all experiments.

### Protein depletion experiments

Anchor-away of FRB-tagged protein was induced by the addition of rapamycin (1 mg/ml of 90% ethanol/10% Tween 20 stock solution) to the culture media to a final concentration of 1 μg/ml for 1h (Haruki et al., 2008). Degradation of aid*-tagged proteins was obtained by addition of IAA to a final concentration of 0.5 μM for 30 min. In experiments in which anchor-away and degron were used simultaneously the cells were treated with rapamycin, after 30 min IAA was added to the culture and cells were grown for another 30 min before harvesting.

The efficiency of protein depletion was monitored by fluorescence microscopy of cells bearing FRB-GFP-tagged fusion proteins. Briefly, cells fixed with cold methanol by a 6-min incubation at -20°C, centrifugated, resuspended in PBS+DAPI solution, incubated for 5 min, washed once and resuspended in PBS for microscopy (Molecular Devices ImageXpress Micro XL). Degradation of aid*-tagged proteins was monitored by western blotting.

### ChEC-seq

ChEC-seq experiments were performed essentially as described (Kubik et al., 2018; Zentner et al., 2015). A strain expressing “free” MNase under the control of the *REB1* promoter was used as a control. Briefly, cells were washed and resuspended in buffer A (15 mM Tris 7.5, 80 mM KCl, 0.1 mM EGTA, 0.2 mM spermine, 0.5 mM spermidine, 1xRoche EDTA-free mini protease inhibitors, 1 mM PMSF) with 0.1% digitonin and incubated for 5 min at 30°C. Calcium chloride was added to the final concentration of 2 mM to induce MNase activity. Reactions were stopped after 1 min by adding EGTA to a final concentration of 50 mM. DNA was purified using MasterPure Yeast DNA purification Kit (Epicentre) and small DNA fragments were preserved by purification with AMPure beads (Agencourt) as described (Kubik et al., 2018). Libraries were prepared using NEBNext kit (New England Biolabs) as described before (Kubik et al., 2018) and sequenced using HiSeq 2500 in single-end mode. Read ends were considered to be MNase cuts and were mapped to the genome (sacCer3 assembly) using bowtie2 through HTSStation (David et al., 2014) and densities of MNase cuts were calculated by extending the 5’ read ends by 1 bp. All densities were normalized to 10M reads.

### MNase-seq

Experiments were performed as described before (Kubik et al., 2018). Yeast cultures were crosslinked, spheroplasted and treated with varying concentrations of MNase for 45 min at 37°C. Reactions were stopped by addition of 30 mM EDTA and the samples were de-crosslinked by overnight incubation at 65°C in the presence of SDS (0.5%) and proteinase K (0.5 mg/ml). DNA was purified by ethanol precipitation, treated by RNase and sequencing libraries were prepared as described (Kubik et al., 2018). The libraries were sequenced using HiSeq 2500 in the paired-end mode. Mapping of the sequencing data to sacCer3 genome assembly was performed using bowtie2 through HTSStation (David et al., 2014). Mapped reads were trimmed by 15 bp from each side when calculating densities to better visualize individual nucleosome peaks. All densities were normalized to 10M reads.

### ChIP-seq

ChIP-seq was performed essentially as described before (Kubik et al., 2018). Crosslinked cells were lysed by bead-beating, chromatin was sonicated, and the soluble fraction was incubated with the appropriate antibody and magnetic beads for 3h. For RNAPII ChIP-seq 5% (v/v) of crosslinked, sonicated *S. pombe* chromatin was added as a spike-in control prior to antibody addition. The beads were washed, and DNA was eluted, de-crosslinked and purified using High Pure PCR Cleanup Micro Kit (Roche). The libraries were prepared using TruSeq ChIP Sample Preparation Kit (Illumina) according to manufacturer’s instructions. The libraries were sequenced using HiSeq 2500 and the reads were mapped to sacCer3 genome assembly using HTSStation (David et al., 2014) (read densities calculated using shift=100 bp, extension=50 bp). All densities were normalized to 10M reads.

### TSS-seq

The experiments were performed as described before (Challal et al., 2018; Malabat et al., 2015). 10% of *S. pombe* cells were added to *S. cerevisiae* cultures to serve as a spike-in control. Total RNA was extracted from the cells using phenol and chloroform and precipitated with ethanol, DNA was digested with DNase I and RNA was extracted and precipitated again. Polyadenylated transcripts were purified using oligo d(T)25 magnetic beads (New England Biolabs). RNA was dephosphorylated using FastAP Thermosensitive Alkaline Phosphatase (ThermoFisher) and treated with Cap-Clip Acid Pyrophosphatase (Tebu-bio). RNA was then ligated to the biotinylated 5’ adaptor and fragmented for 5 min at 70°C in fragmentation buffer (10mM ZnCl2, 10mM Tris pH7.5). The reaction was stopped with 1 µl of 0.5 M EDTA. Ligated RNA molecules were purified using streptavidin magnetic beads (New England Biolabs). Reverse transcription was performed with RevertAid reverse transcriptase (ThermoFisher) and cDNAs were purified with Agencourt RNAClean XP beads (Beckman Coulter). DNA was amplified with LA Taq DNA polymerase (Takara) and purified with NucleoMag NGS Clean-up and Size Select (Macherey-Nagel). The resulting libraries were sequenced in single-end mode and the results were mapped to sacCer3 genome assembly.

### ChEC-seq signal normalization

In our previous work (Kubik et al., 2018), ChEC-seq was normalized by calculating the ratio between the ChEC-seq tag counts at a position (i.e. Rsc8-MNase cuts sites) and the tag counts of free MNase at the same site. Although this approach is generally correct and robust for Rsc8 it has a serious disadvantage: regions of low cut frequency tend to have high variation in signal ratio that might not reflect true binding events but instead result from a random fluctuation of the sequencing signal. This increases the noise in the data and reduces the possibility of finding true binding events, particularly for weak sites. Smoothing the ratio by calculating an average of ratio values in neighboring sites partially reduces the noise but at the same time reduces the precision of the technique.

We turned to a non-parametric normalization method for ChEC-seq data that reduces the noise without reducing the precision. Our method uses an empirical Bayesian estimation of the prior distribution (in this case, the ratio between the ChEC-seq signal for the tested protein and for free-MNase) to increase the signal to noise ratio by reducing the effect of random fluctuations in low coverage areas. As a result, at low-coverage regions the ratio is decreased to the genome-wide average. Empirical Bayes estimation uses signal ratios (scaled between 0 and 1) as the prior. Distribution of the ratios calculated genome-wide is fit to the beta distribution and the α and β parameters of the distribution are used to adjust the signal ratio according to the equation 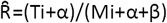, where Ti is the signal in the test sample and Mi is the signal for control (free MNase). Such adjusted ratio was used in all subsequent analysis of ChEC signal.

### ChEC peak calling and clustering

Peaks of protein binding signal were determined from genome-wide normalized ChEC ratio (see above) using Peak locator (Kubik et al., 2015) with the minimal normalized signal threshold of 5 and the window size of 100 bp. Peaks determined for different remodelers were pooled and all regions found within 150 bp of each other were merged. This common list of all chromatin remodeler binding sites was used to calculate the average normalized signal for each remodeler +/-75 bp from the midpoint of each region. For analysis of promoters, signal was calculated in the region spanning -250 to -100 bp from the dyad of the +1 nucleosome for every gene with a well annotated TSS (van Bakel et al., 2013). In the next step, each region displaying a signal of at least 2 was assigned the value of 1 and below 2 was assigned the value of 0. For Swi3, due to significantly higher peak signals, the threshold was set to 6. The list was then k-means clustered according to the 0/1 values with k=8, excluding data for remodelers whose depletion did not significantly affect promoter nucleosomes (i.e. ISW1 and CHD1).

### Nucleosome occupancy and stability change

Nucleosome occupancy change (either positive or negative) was calculated in 10 bp windows as the log2 ratio of read counts in remodeler-depleted cells compared to mock-treated cells (using high concentration MNase-seq data; **Figure 4D**). To quantify the average overall magnitude of nucleosome occupancy change at promoter regions, absolute values for read count differences between CR-depleted and mock-treated cells were used (**Figures 2C, 3B, 3D**).

Nucleosome changes related to +1 nucleosome were considered significant if the absolute log2 ratio of occupancy, calculated in the region spanning -/+ 150 bp from +1 dyad, was higher than 0.7/bp. To estimate nucleosome stability changes nucleosome occupancy was calculated in a region -/+50 bp from each nucleosome dyad in remodeler-depleted and mock-treated cells. A fragile nucleosome was considered to become stabilized by remodeler depletion if its average occupancy in the high MNase assay increased by at least 15 reads and its dyad was found within 50 bp of its original position (from (Kubik et al., 2015)). A nucleosome was considered to be destabilized by remodeler depletion if its average occupancy in the high MNase assay decreased by at least 15 reads and its dyad was not found within 73 bp of its original position (determined in mock-treated cells).

### ChIP-seq spike-in normalization and quantification

RNAPII ChIP-seq signal in *S. cerevisiae* was normalized using a *S. pombe* spike-in control as described before (Bruzzone et al., 2018). TBP binding was calculated in regions spanning 200 bp centered on all TATA and TATA-like sites, taken from (Rhee and Pugh, 2012). RNAPII binding signal was calculated inthe transcribed region of all genes with well determined TSSs and TTSs (based on (van Bakel et al., 2013))and in the ORF for all other genes. Changes in RNAPII signal were considered significant if the RNAPII decreased/increased by 1.5-fold and the average signal was at least 30 reads/bp in mock-treated or remodeler-depleted samples for up-regulated and down-regulated genes, respectively. Genes were considered as not affected if the log2 change in RNAPII signal was in the range >-0.1 and <0.1 and the average signal in the mock-treated sample was at least 30 reads/bp.

### TSS determination

TSS signals from three replicates of each experiment were averaged separately for the Watson and the Crick strand. For the analysis shown in Figure 7 all TSSs in “puller”-depleted cells were found with a minimum signal of 150 reads. For each peak the nearest ATG on the respective strand was found (at a maximum distance of 500 bp) and then a single, strongest TSS was identified for each gene. Signal was calculated for each of these TSSs in the wild-type and “puller”-depleted conditions. A significant decrease or increase in signal was set at a threshold of log2ratio = +/-1. Regions displaying artefactually high signal (e.g. found near rDNA) were removed from the analysis.

### TATA-box search

All putative TATA-box sites were searched for by first looking for matches to the canonical TATAWAWR motif using FIMO from the MEME Suite with a threshold of p<0.001. Searches were also performed for motifs with up to 2 substitutions in the consensus or using the frequency matrix determined for TBP (Spt15) binding (Khan et al., 2018).

All three types of searches yielded very similar motif frequencies for both up- and down-regulated gene classes shown in **Figure 7**.

### Plots and statistics

Figures 1C, 4D, S1B, S1C, S5A, S5F and S7 were made using EaSeq (Lerdrup et al., 2016). For box-and-whisker plots the following elements indicate: center line, median; box limits, upper and lower quartiles; whiskers, 1.5x interquartile range. Statistic test were applied where indicated.

### Data and software availability

All sequencing data generated in this study were submitted to the GEO database as Series GSE115412 (for ChEC-seq, MNase-seq and ChIP-seq) and Series GSE114589 (TSS-seq).

## Supplemental Figure legends

**Figure S1.**
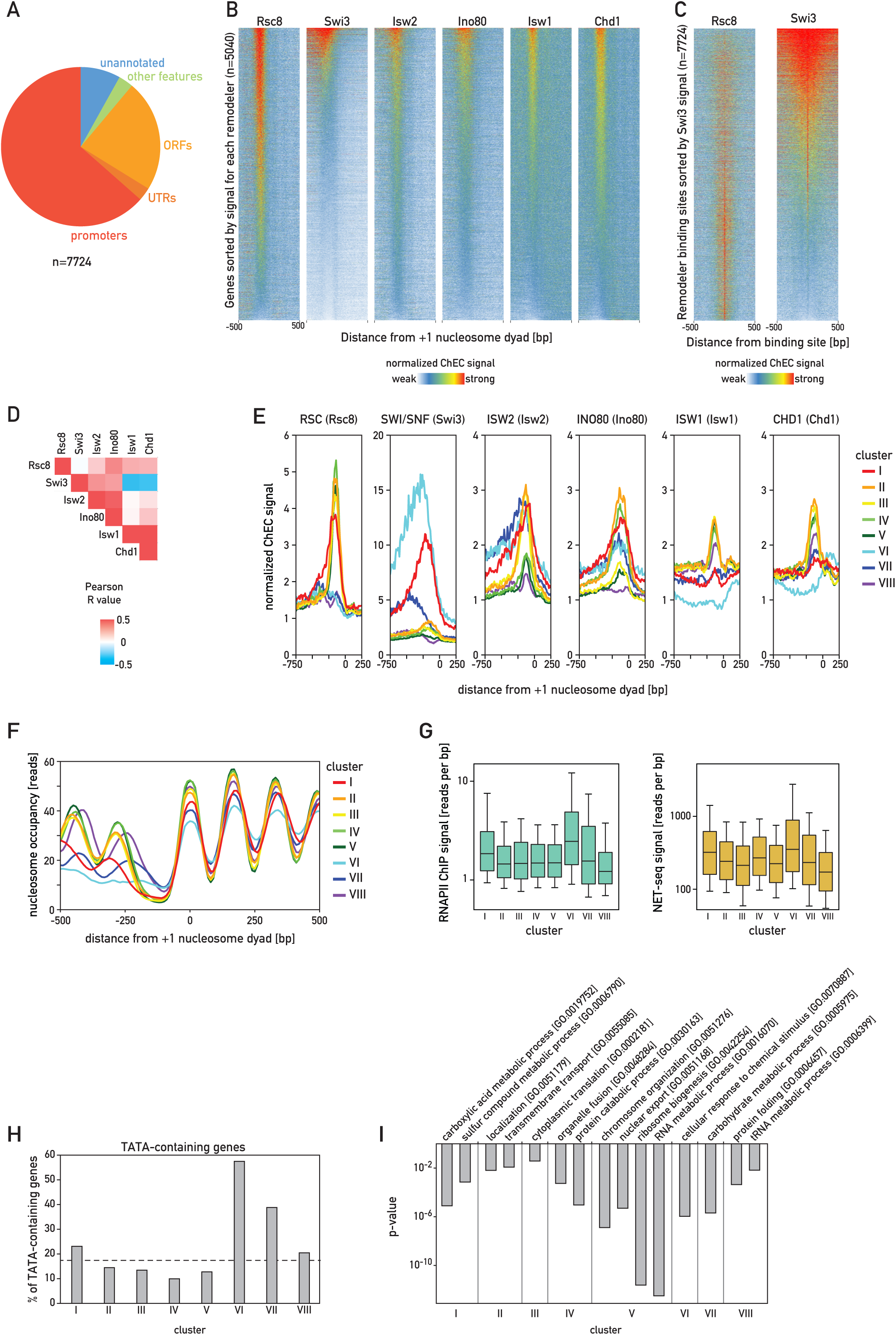
(**A**) Distribution of identified remodeler binding sites between different genomic features. (**B**) Heatmaps displaying ChEC signal for every CR centered on +1 nucleosome of 5040 protein-coding genes and sorted individually by the signal of each remodeler calculated in the promoter (-250 to -50 bp from +1 dyad). (**C**) Heatmaps displaying ChEC signal for Rsc8 and Swi3 at all sites displaying remodeler binding sorted by the signal for Swi3. (**D**) Grid representing Pearson correlation coefficients between ChEC signal for different CRs calculated at all identified remodeler binding sites. (**E**) Plots displaying average normalized ChEC signal for each CR calculated for all clusters. (**F**) Average nucleosome occupancy in each cluster. (**G**) Boxplots displaying expression of genes belonging to each cluster measured either as RNAPII ChIP-seq signal in the gene body or NET-seq signal (Churchman et al., 2011). (**H**) Fraction of genes in each cluster which contain a canonical TATA-box (according to Rhee et al., 2012), dashed line indicates genomic average (∼17%). (**I**) Chosen, enriched GO-terms representative for each cluster.

**Figure S2.**
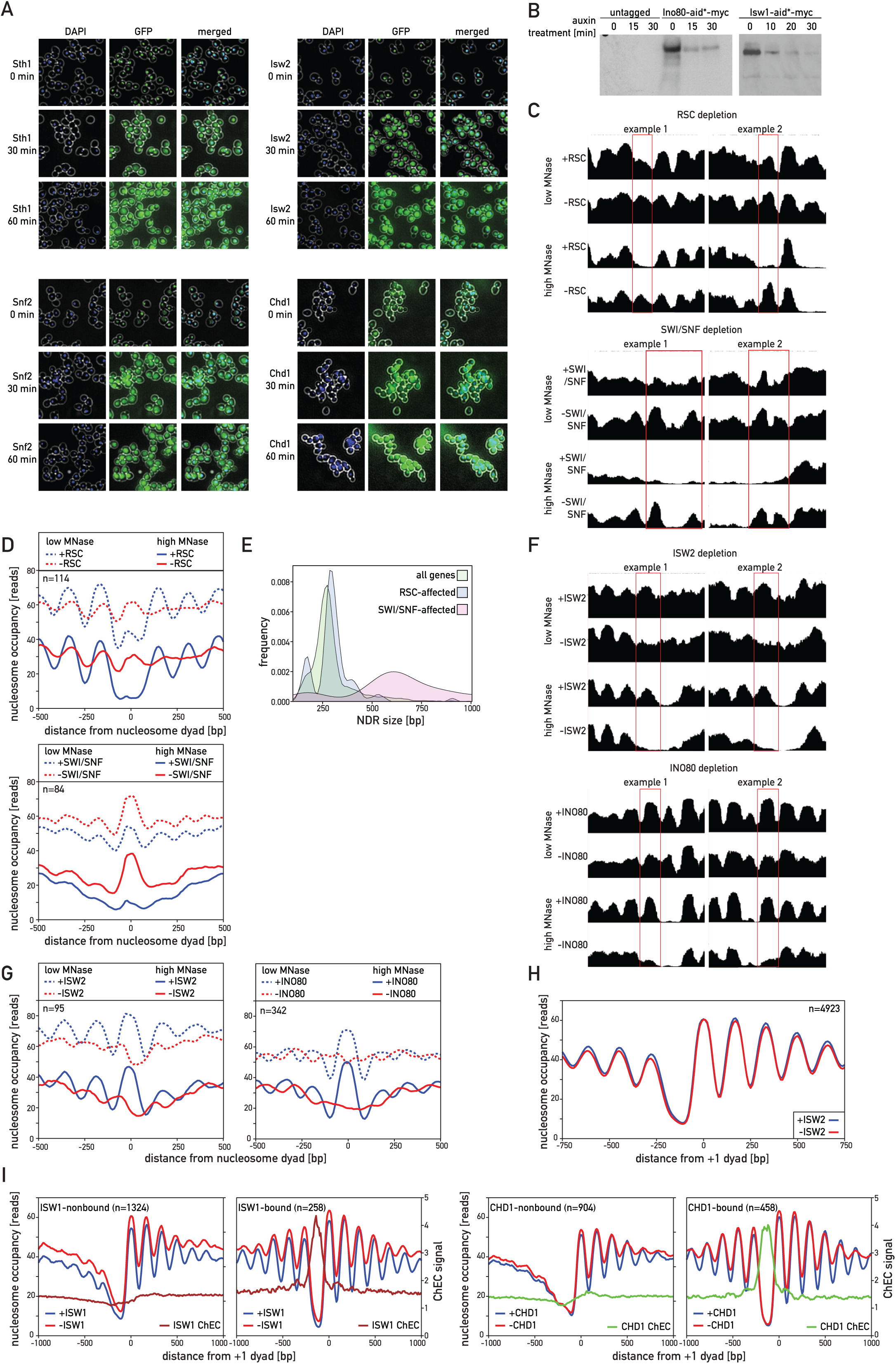
(**A**) Fluorescence microscopy of cells bearing FRB-GFP fusions of Sth1, Snf2, Isw2 and Chd1; cells were treated with rapamycin for indicated times, fixed and stained with DAPI. (**B**) Western blotting (anti-myc antibodies) of cell lysates from an untagged strain and strains bearing Ino80-aid*-myc and Isw1-aid*-myc fusions treated with auxin for indicated amount of time. (**C**) Snapshots of sample regions in which nucleosomes become stabilized (marked with a red rectangle) upon depletion of RSC (top) or SWI/SNF (bottom). (**D**) Average plots of nucleosome occupancy for all nucleosomes becoming stabilized upon depletion of RSC (top) or SWI/SNF (bottom). (**E**) Frequency of genes according to their NDR size for genes with promoter nucleosome stability decreased upon RSC depletion, SWI/SNF depletion or for all genes. (**F**) Snapshots of sample regions in which nucleosomes become destabilized (marked with a red rectangle) upon depletion of ISW2 (top) or INO80 (bottom). (**G**) Average plots of nucleosome occupancy for all nucleosomes becoming destabilized upon depletion of ISW2 (left) or INO80 (right). (**H**) Average plot of nucleosome occupancy upon depletion of ISW2 for all genes with unaffected +1 nucleosome occupancy. (**I**) Nucleosome occupancy and ChEC signal at genes with lowest (left) or highest (right) binding of ISW1 or CHD1.

**Figure S3.**
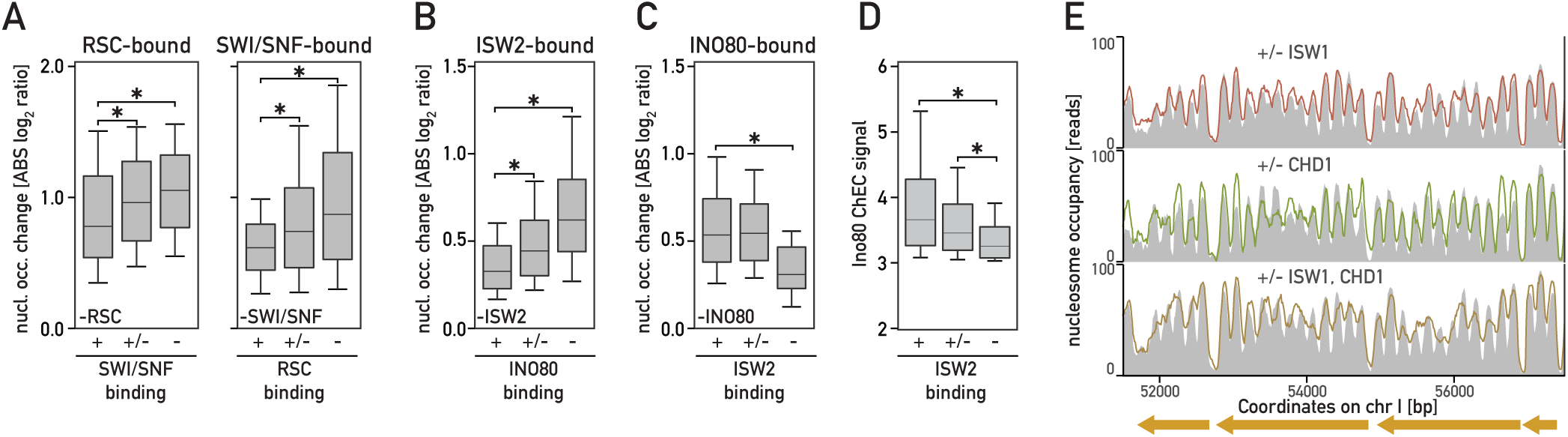
(**A**) Nucleosome occupancy change upon depletion of RSC (left) or SWI/SNF (right) at sites bound by each remodeler and displaying varying binding signal of the other one. (**B**) Nucleosome occupancy change upon depletion of ISW2 at sites bound by this remodeler and displaying varying binding signal of INO80. (**C**) Nucleosome occupancy change upon depletion of INO80 at sites bound by this remodeler and displaying varying binding signal of ISW2. (**D**) Boxplot of INO80 binding signal at INO80-bound sites displaying varying binding signal of ISW2. **(A-D)** Asterisk indicates significant difference (p<0.05, Mann-Whitney test). (**E**) Snapshot of a sample genomic region displaying nucleosome occupancy change upon depletion of ISW1, CHD1 or both remodelers simultaneously.

**Figure S4.**
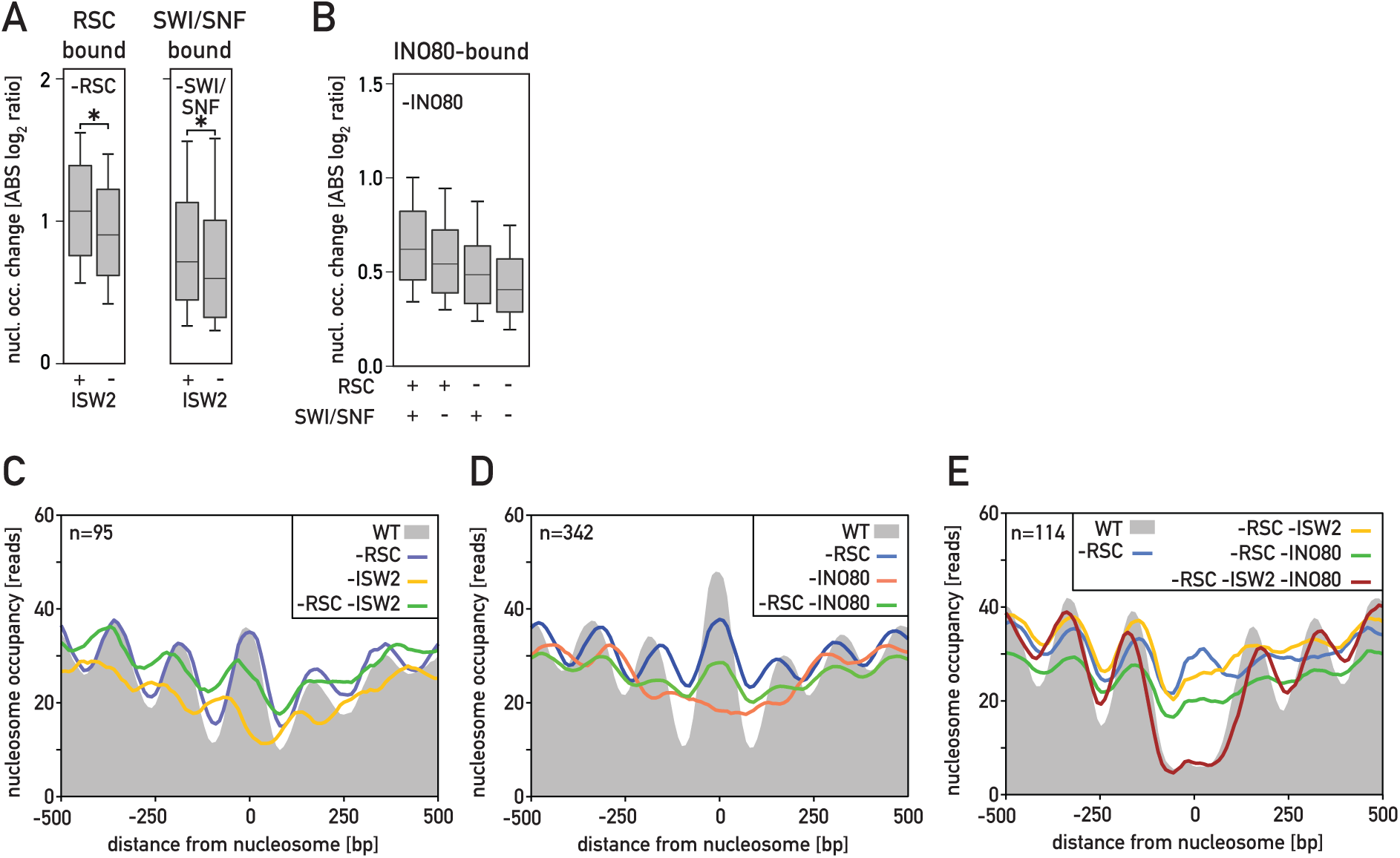
(**A**) Nucleosome occupancy change upon RSC depletion at sites bound by RSC (left) and change upon SWI/SNF depletion at sites bound by SWI/SNF (right) for sites co-bound by ISW2 or not. Asterisk indicates significant difference (p<0.05, Mann-Whitney test) (**B**) Nucleosome occupancy change upon INO80 depletion at sites bound by INO80 and co-bound by RSC and/or SWI/SNF or not. (**C**) Average plots of nucleosome occupancy for all nucleosomes becoming destabilized upon depletion of ISW2 shown for wild-type cells and cells depleted of RSC, ISW2, or both remodelers simultaneously. (**D**) Average plots of nucleosome occupancy for all nucleosomes becoming destabilized upon depletion of INO80 shown for wild-type cells and cells depleted of RSC, INO80, or both remodelers simultaneously. (**E**) Average plots of nucleosome occupancy for all nucleosomes becoming stabilized upon depletion of RSC shown for wild-type cells and cells depleted of RSC, RSC and ISW2, RSC and INO80, or all three remodelers simultaneously.

**Figure S5.**
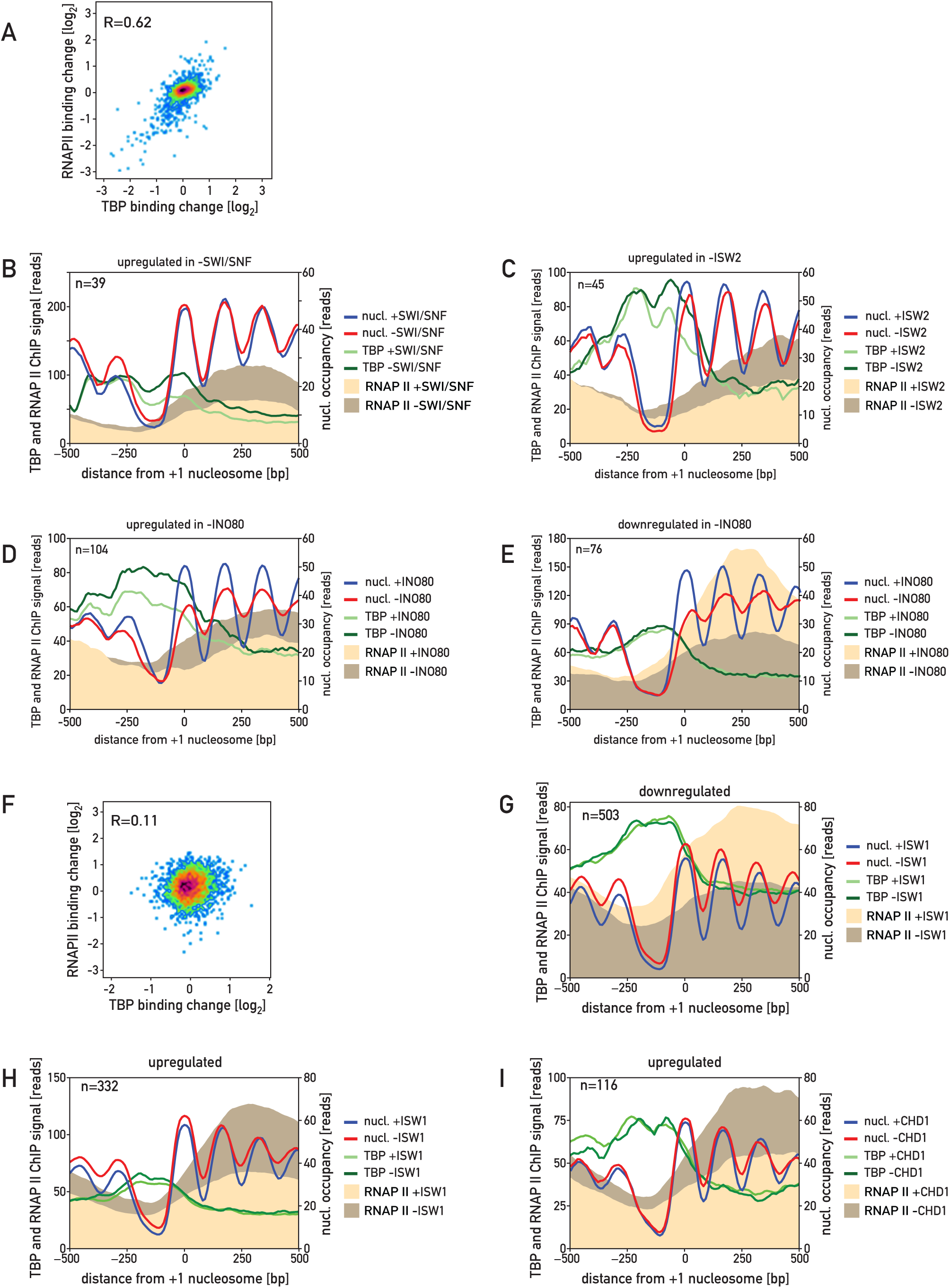
(**A**) Scatterplot showing relationship between TBP binding change at gene promoters and RNAPII binding change in corresponding gene bodies upon SWI/SNF depletion. (**B**) Plots displaying nucleosome occupancy, RNAPII and TBP ChIP signals, in the presence and absence of SWI/SNF, at genes displaying a significant increase in RNAPII level upon SWI/SNF depletion. (**C**) Plots displaying nucleosome occupancy, RNAPII and TBP ChIP signals, in the presence and absence of ISW2, at genes displaying a significant increase in RNAPII level upon ISW2 depletion. (**D**) Plots displaying nucleosome occupancy, RNAP II and TBP ChIP signals, in the presence and absence of INO80, at genes displaying a significant increase in RNAPII level upon INO80 depletion. (**E**) Same as in (D) for downregulated genes. (**F**) Scatterplot of TBP signal change and RNAPII signal change at corresponding genes. (**G**) Plots displaying nucleosome occupancy, RNAPII and TBP ChIP signals, in the presence and absence of ISW1, at genes displaying a significant decrease in RNAPII level upon ISW1 depletion. (**H**) As in (G) but for upregulated genes. (**I**) Plots displaying nucleosome occupancy, RNAPII and TBP ChIP signals, in the presence and absence of CHD1, at genes displaying a significant increase in RNAPII level upon CHD1 depletion.

**Figure S6.**
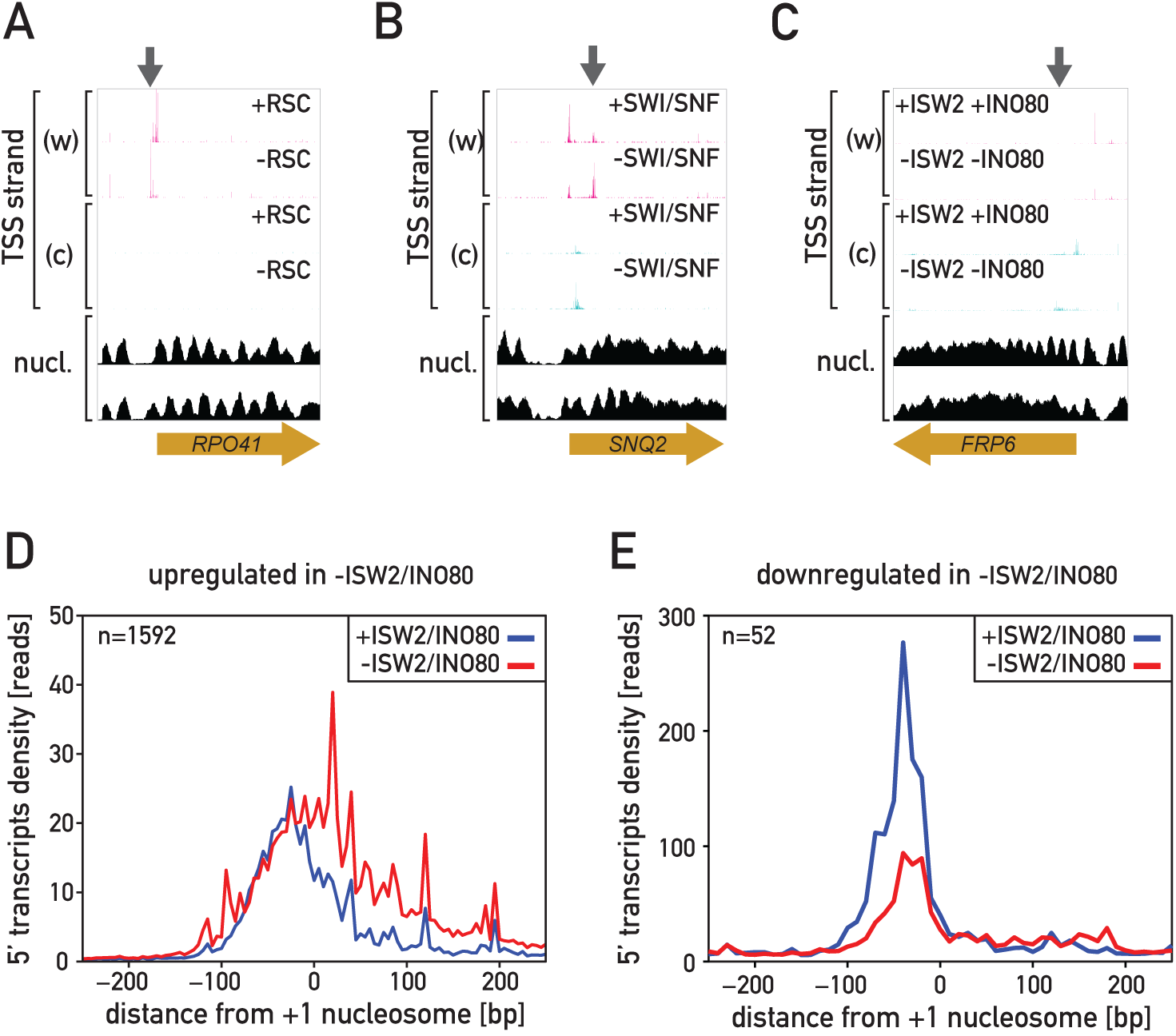
(**A**) Snapshot of genomic region showing 5’RACE signal for the Watson (w) and the Crick (c) strands as well as nucleosome occupancy in the presence and absence of RSC; upon RSC depletion the *RPO49* gene transcription initiates more upstream comparing to wild-type conditions. (**B**) As in (A) but for SWI/SNF depletion; upon SWI/SNF depletion the *SNQ2* gene transcription initiates more frequently downstream comparing to wild-type conditions; additionally, there is more initiation events in the opposite strand just downstream from the upstream-most *SNQ2* TSS. (**C**) As in (A) but for ISW2 and INO80 simultaneous depletion; upon “pullers” depletion the *FRP6* gene transcription initiates at a downstream position comparing to wild-type conditions. (**D**) Average 5’RACE signal at genes upregulated upon simultaneous depletion of ISW2 and INO80. (**E**) As in (D) but for downregulated genes.

**Figure S7.**
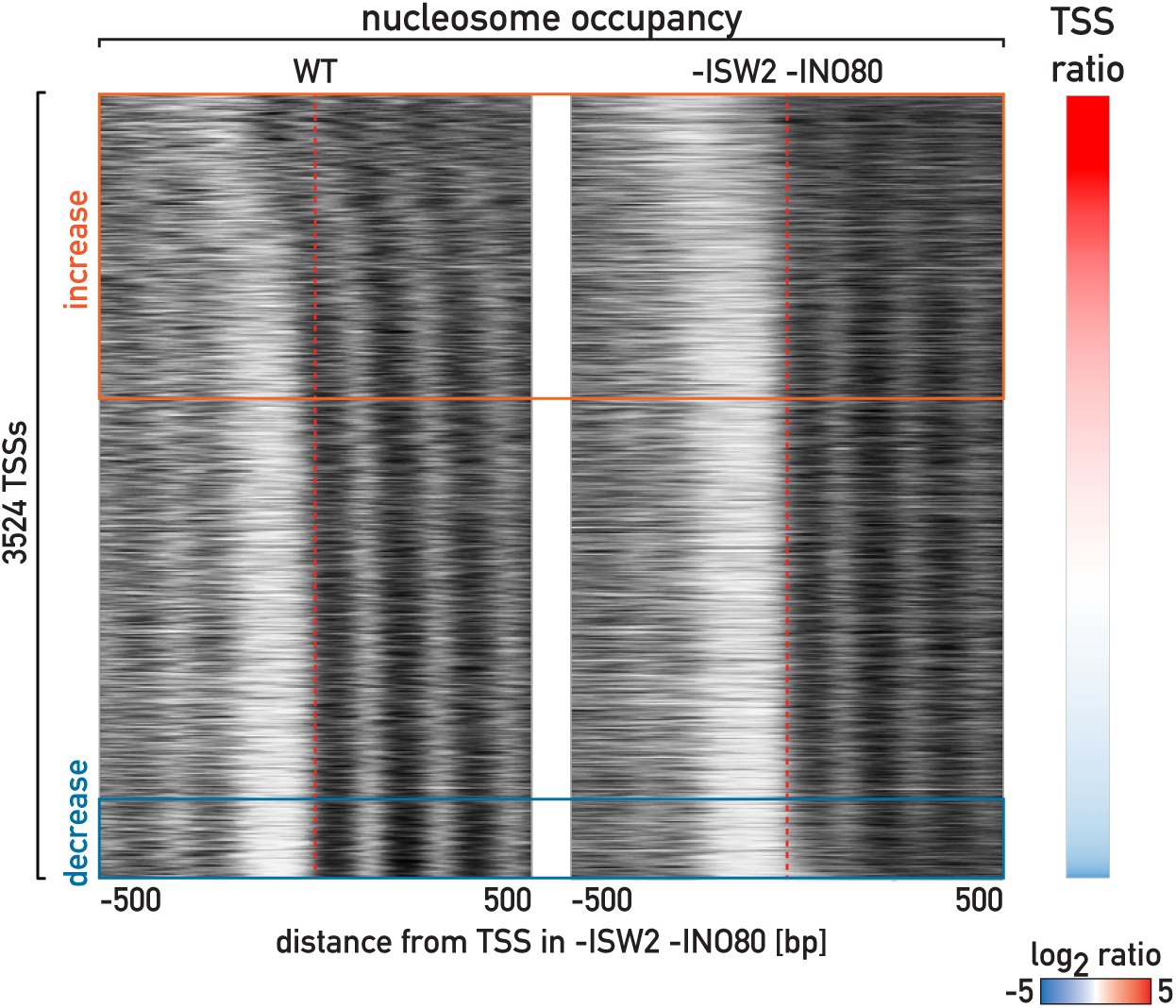
Heatmaps showing nucleosome occupancy, centered at TSS positions determined in cells depleted of ISW2 and INO80, shown in wild type cells and cells depleted of both remodelers.

